# Elements of divergence in germline determination in closely related species

**DOI:** 10.1101/2022.08.12.503758

**Authors:** Shumpei Morita, Nathalie Oulhen, Stephany Foster, Gary M. Wessel

## Abstract

Evolutionary transitions enable the wide diversity in life histories of plants and animals. This is particularly germane in the development of the germ line in which fitness is a direct readout of evolutionary change. Here, we focused on the gem line of two distinct sea urchin species who shared a common ancestor 50 million years ago. Even though they both rely on inherited mechanisms to specify their germ line, the integration of stage-matched single cell RNA-seq (scRNA-seq) datasets from these two sea urchins revealed a variety of differences in gene expression, including a broader expression of the germ line factor Nanos2 in *Lytechinus variegatus* (*Lv*) compared to *Strongylocentrotus purpuratus* (*Sp*). In *Sp, Nanos2* mRNA expression is highly restricted to the primordial germ cells (PGCs) by a lability element in its 3’UTR. This element is lacking in the mRNA of *Lv Nanos2*, explaining how this mRNA more broadly accumulates in the *Lv* embryos. We discovered that the *Lv Nanos2* 3’UTR instead leads to a germline specific translation of the protein. The results emphasize that regulatory mechanisms resulting in germline diversity rely less on transcriptional regulation and more on post-transcriptional and post-translational restrictions of key gene products, such as Nanos2.

**Highlights:**

- The first integration of scRNA-seq datasets comparing two echinoderm species.
- We find Nanos2 positive cells in the embryonic soma of *Lytechinus variegatus*, an unusual occurrence, but not in *Strongylocentrous purpuratus*.
- We discovered that this somatic *Nanos2* mRNA is lacking an important regulatory element (GNARLE) in its 3’UTR
- Instead, in *Lv*, the 3’UTR of *Nanos2* leads to its specific translation in the germ cells.

## Introduction

The germ line is one of the few cell lineages shared by nearly all animals; it generates the eggs and sperm necessary for sexual reproduction. However, the germ line is formed by highly diverse mechanisms, sometimes even between closely related taxa. The marked variations in these mechanisms is likely a node of evolutionary manipulation, and selected for by success in fitness. Such a direct impact on fitness is likely a common feature of germ cell diversification, but how such transitions in mechanism have occurred is only beginning to be appreciated.

The primordial germ cells (PGCs) are specified during embryogenesis to give rise to the germ line; Nanos is an RNA binding protein essential for the maintenance and survival of the PGCs. Its function has been tested in multiple species (Kobayashi *et al*., 1996, Subramaniam and Seydoux, 1999, Koprunner *et al*., 2001, Tsuda *et al*., 2003, Juliano *et al*., 2010). Together with its partner Pumilio, Nanos binds to a specific element in the 3’UTR of its target mRNAs, the PRE (pumilio response element). This binding induces the translational inhibition and/or the degradation of the targeted mRNAs. So far, only a few targets of the Nanos/Pumilio complex have been identified: hunchback, cyclin B, hid, VegT, CNOT6, eEF1a (Dalby and Glover, 1993, Murata and Wharton, 1995, Wreden *et al*., 1997, Asaoka-Taguchi *et al*., 1999, Kadyrova *et al*., 2007, Sato *et al*., 2007, Lai *et al*., 2012, Swartz *et al*., 2014, Oulhen *et al*., 2017). Nanos is rigidly regulated and its ectopic expression leads to embryonic lethality (Luo *et al*., 2011). However, a variety of methods are employed by animals to restrict Nanos to the PGCs.

In the sea urchin, the PGCs arise from two asymmetric cell divisions, resulting in the four small micromeres at the 5th cell division (32 cell stage). Shortly after their formation, their cell cycle is downregulated, they divide only once more by the end of gastrulation, and they show a transient downregulation of their transcriptional, translational, RNA degradation and mitochondrial activities (Oulhen *et al*., 2017, Oulhen and Wessel, 2017). In the sea urchin *Strongylocentrotus purpuratus* (*Sp*), Nanos2 is essential to maintain this transient quiescence. Multiple levels of regulation restrict *Sp Nanos2* mRNA and protein to the PGCs early in development: it is transcribed broadly in the early embryo through the Wnt pathway (Oulhen *et al*., 2019b, Pieplow *et al*., 2021) but its mRNA only accumulates in the PGCs. *Sp Nanos2* contains an element (GNARLE) in its 3’UTR that leads to its degradation in the somatic cells and its retention in the PGCs (Oulhen *et al*., 2013). And finally, it is not possible to overexpress this protein in the somatic cells, since the protein itself possesses regulatory elements that leads to its degradation in somatic cells and its retention in the PGCs (Oulhen and Wessel, 2016).

More recently, we used single-cell RNA-seq (scRNA-seq) to identify the transcriptome of the PGCs throughout the development of *Sp* embryos. As expected, the transcript coding for Nanos2 was highly restricted to the PGC cluster (Foster *et al*., 2020). Here we use scRNA-seq analysis to compare the developmental profile of two sea urchin species with a last common ancestor of ∼50 myr, *S. purpuratus* (*Sp*) and *L. variegatus* (*Lv*). Although both species appear to specify their PGCs by inherited mechanisms, they are highly divergent in detail. This scRNA-seq analysis suggests that Nanos2 expression is one of the most differently regulated genes between these sea urchin species. *Sp* Nanos2 mRNA is tightly restricted to the PGCs, whereas *Lv Nanos2* mRNA is more broadly expressed throughout the embryo. We show that this accumulation is a result of the RNA-lability element present in the 3’ UTR of *Sp*, but not in *Lv*.

## Results

### Single-cell RNA-seq analysis of *Lv* embryos

A previous study reported the scRNA-seq analysis of fixed cells from sea urchin embryos of *Lytechinus variegatus* (Massri *et al*., 2021); the authors analyzed 18 timepoints across the first 24 hours of development, capturing 50,935 total cells. Here, for the purpose of integrating and comparing scRNA-seq datasets from *Strongylocentrotus purpuratus* and *Lytechinus variegatus*, we generated an *Lv* scRNA-seq dataset using live cells with the same protocol and morphological stages as described for *Sp (*Foster *et al*., 2020). We cultured *Lv* embryos and processed eight time points from two hours post-fertilization (2H) to late gastrula stage (Fig.1A). In total, the transcriptome profiles of 63,930 cells were analyzed in this dataset (Table S1). The datasets of the eight time points were integrated into a single dataset (*Lv* dataset) and cells were grouped into clusters by Uniform Manifold Approximation and Projection (UMAP) (Becht *et al*., 2018, McInnes, 2018) (Fig.S1A). However, the cells were clustered by developmental time points rather than by transcriptional profiles probably due to technical batch-effects among samples (Fig. S1A and B). To overcome this issue, we performed batch-effect correction to align all developmental time points by using Harmony (Fig. S1C) (Korsunsky *et al*., 2019). In the Harmony-normalized *Lv* dataset, cell populations derived from the eight developmental time points overl(Massri *et al*., 2021)apped each other with high fidelity (Fig. S1C and D). This result indicates that the batch-effect correction procedure enabled us to compare transcriptional profiles between stages and characterize cell types throughout embryogenesis.

**Fig.1.**
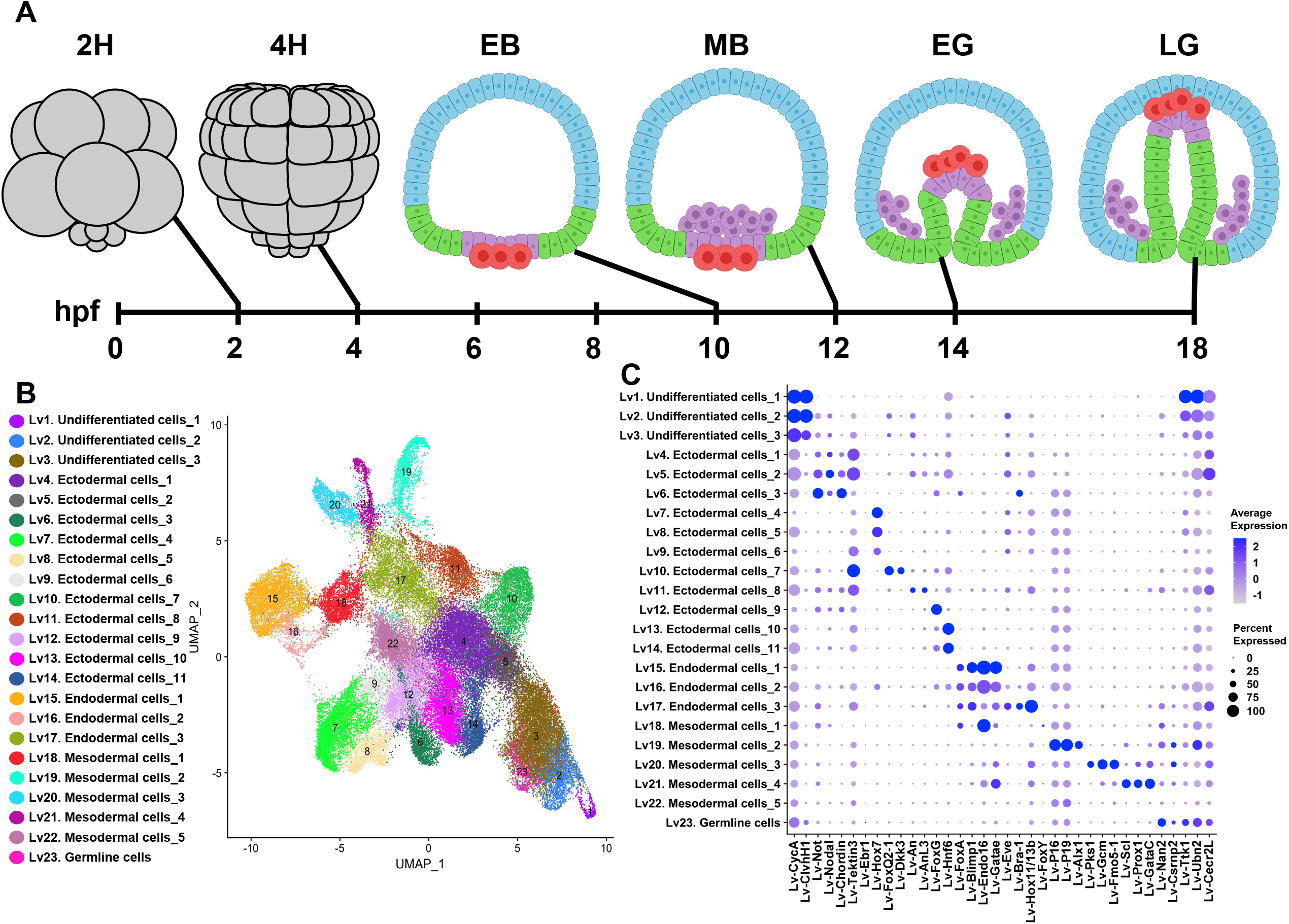
Identification of 23 cell-populations in *Lytechinus variegatus* embryos. **(A)** Diagram of sea urchin embryogenesis. We collected embryos at eight developmental time points from 2 hours post-fertilization (hpf) to 18 hpf. The diagrams of embryos at 2 hpf (2H), 6 hpf (2H), 10 hpf (EB), 12 hpf (MB), 14 hpf (EG) and 18 hpf (Tischler *et al*.) are illustrated. Undifferentiated cells (gray), ectodermal cells (blue), endodermal cells (green), mesodermal cells (purple) and germline cells (Chatfield *et al*.) are shown. This figure was created with BioRender.com (https://app.biorender.com/) **(B)** Identification of 23 cell-populations using UMAP visualization for integrated dataset of 8 developmental time points. **(C)** Dot plot represents marker gene expression characterizing cell types in the *Lv* dataset. Dot size and dot color indicate the percentage of cells expressing the gene and the average expression level, respectively.

In our *Lv*-dataset, we identified 23 cell states (Fig. 1B and C). Among them, three cell states (Lv1–Lv3) were identified as undifferentiated cells because these were predominantly observed in the cleavage stages (2H–8H) and did not show characteristic marker gene expression as described below (Fig. S2). We also identified eleven ectodermal clusters (Lv4-Lv14), three endodermal clusters (Cluster Lv15–Lv17), and five mesodermal cell states (Cluster Lv18– Lv22). These cell states were identified according to the expression of well-described markers previously published (Angerer *et al*., 1989, Ettensohn *et al*., 2003, Otim *et al*., 2004, Cheers and Ettensohn, 2005, Duboc *et al*., 2005, Angerer *et al*., 2006, Oliveri *et al*., 2006, Tu *et al*., 2006, Massri *et al*., 2021). Markers for each of these clusters are presented in Table S2.

A previous single cell RNA seq dataset was published for this animal, 18 time points throughout embryonic development were analyzed to explore cell trajectories using the Waddington-OT (Massri *et al*., 2021). Even though the normalization, the method and the goals were different from the dataset presented here, similar cluster markers were obtained. For example, as expected, both datasets showed an enrichment of Alx1, Gcm, Pks1, Scl in the mesodermal cell states, Endo16, FoxA and Blimp in the endodermal cell states, Chordin, Nodal, FoxQ2_1 in the ectodermal cell states.

### Transcriptional profile of the PGCs in *Lv* embryos

Our previous study revealed that *Lv Nanos2* is transcribed during early embryogenesis and first accumulated in cells in the vegetal region at the 128-cell stage (Fresques *et al*., 2016). In this study, we employed scRNA-seq analysis to investigate *Lv Nanos2* expression during embryogenesis. Consistent with the previous study, no or low expression of *Lv Nanos2* was detected at 2H and 4H post-fertilization but it is then upregulated at 6H and 8H post-fertilization (Fig.2A–D). However, the expression was not enriched in a specific cell population (Fig.2C and D). Notably, in early (EB) and mesenchyme (MB) blastula, endodermal and mesodermal cell populations (Lv16–Lv22) tended to express higher levels of *Lv Nanos2* mRNA compared to ectodermal cell populations (Lv4–Lv15) (Fig. 2E and F), suggesting that *Lv Nanos2* mRNA is expressed primarily in the vegetal plate, which is composed of endo-mesodermal cells and PGCs (Fig. 1A). *Lv Nanos2* expression gradually became prominent in Lv23 from the EB stage onward (Fig. 1C and 2E–H), Lv23 was identified as the germline. Somatic cells though, exhibited significant *Lv Nanos2* expression, especially the undifferentiated cells (Lv2 and Lv3), animal ectoderm (Lv10 and Lv11) and mesodermal cells (Lv18 and Lv21) in the early and late gastrula stages (Fig. 2G and 2H).

**Fig.2.**
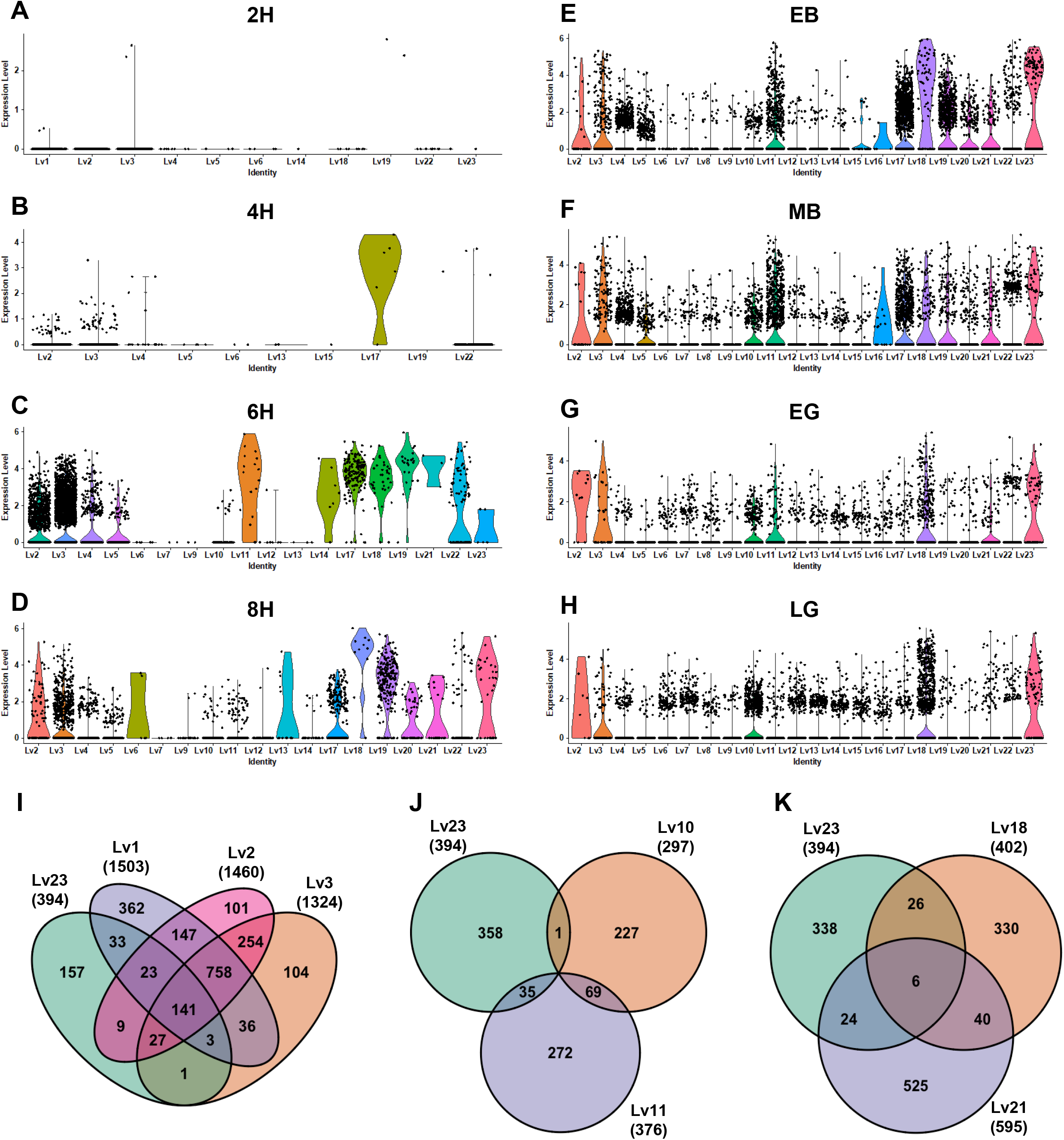
Transcriptional profile of Nanos2-expressing cells in *Lv* embryos. (A–H) Violin plots showing *Lv Nanos2* expression in 2H (A), 4H (B), 6H (C), 8H (D), EB (E), MB (F), EG (G) and LG (H) embryo. Dots represent expression level of Nanos2 in individual cells. (I–K) Venn diagrams showing the number of shared marker gene between germline cells and undifferentiated cells (I), ectodermal cells (J) and endo/mesodermal cells (K).

Since *Nanos2* is one of the most extensively studied genes governing germline development, we hypothesized that the *Nanos2* expressing somatic cells have overlapped marker gene expression with the germ cells. To test this hypothesis, we analyzed how many marker genes are shared between germ cells and *Nanos2*-expressing somatic cells. First, germ cells were compared with undifferentiated cell populations (Lv1–Lv3). We identified 394 marker genes enriched in the germ cells (Table S2). Among them, 141 genes (36%) were shared with all of three undifferentiated cell clusters, and 237 genes (60%) were shared with at least one cluster (Fig. 2I). Next, the germ cells were compared with *Nanos2*-expressing ectodermal cell populations (Lv10 and Lv11) and mesodermal cell populations (Lv18 and Lv21). In contrast to the undifferentiated cells, none, or as few as six marker genes were shared among ectodermal cell and mesodermal cell populations, respectively (Fig. 2J and K). These results show that, even though *Lv Nanos2* expression was observed in several somatic cell populations, the transcriptional profiles were clearly different from that of the germ cells. The transcriptional profile of the germ cells instead was more similar to that of the undifferentiated cell populations containing abundant maternal transcripts (Fig. 2I).

### Three PGC subclusters were identified in *Lv* embryos

The germ cell cluster Lv23 can be subclustered into three subpopulations (Germline_Lv1–3) (Fig. 3A and A’). Germline_Lv1 appeared in the 6H embryos as the sole PGC population but its proportion eventually decreased as the embryo developed (Fig. 3A’ and B). Germline_Lv2 emerged at 8H post-fertilization (Fig. 3A’ and B), its proportion peaked at mesenchyme blastula and decreased by about 7% at the late gastrula stage. The third subpopulation was first detected in the early blastula embryo. Although the proportion was less than 7% in both the EB and MB stages, it was elevated drastically by about 60% at the EG stage, when the proportion of Germline_Lv1 was drastically decreased (Fig. 3B). The proportion of Germline_Lv3 reached over 90% at the LG stage (Fig. 3B).

**Fig.3.**
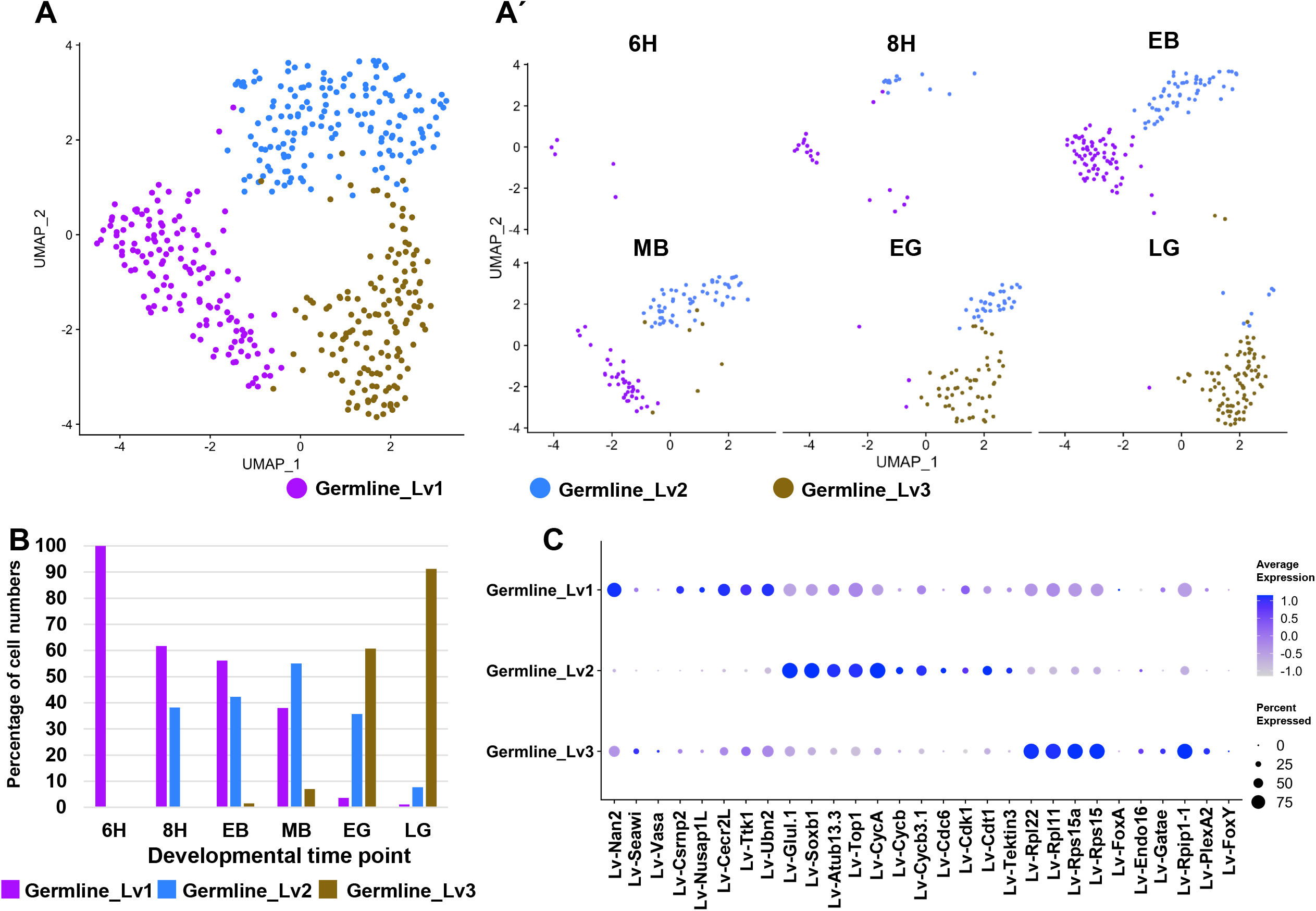
Gene expression changes during germline development in *Lv* embryos. (A and A’) UMAP plot containing the germ cells (A) and UMAP plots split by developmental stages (A’) are presented. (B) The proportion of germline subpopulations in each developmental stage. (C) Dot plot showing marker gene expression in the germline subpopulations. Dot size and dot color indicate the percentage of cells expressing the gene and the average expression level, respectively.

We then analyzed differential gene expression among the three germline subpopulations. We tested well-conserved germline genes (*Nanos2, Seawi* and *Vasa*) and germline-enriched genes (*Lv-Ttk1, Lv-Cecr2L, Lv-Csrnp2, Lv-Nusap1L* and *Lv-Ubn2*), which were identified as the marker genes of the germline cell population (Lv23) in this *Lv*-dataset (Fig. 1C). Almost all of these genes except for Seawi and Vasa were significantly enriched in Germline_Lv1 (Fig. 3C, Fig. S3 and Table S7). By contrast, Germline_Lv2 did not show significant germline gene expression (Fig. 3C, Fig. S3 and Table S7). The top-5 marker genes of Germline_Lv2 included *Lv-Glul, Lv-Soxb1, Lv-Atub13, Lv-Top1* and *Lv-CycA*, all of which were most likely to be maternally deposited transcripts, because of its enriched expression in undifferentiated cell populations (Fig. 1C and 3C; Table S2 and S7). Further, cell cycle-related genes including *Lv-CycA, Lv-CycB, Lv-CycB3, Lv-Cdc6, Lv-Cdk1* and *Lv-Cdt1* were highly expressed in Germline_Lv2 (Fig. 3C). Although germline genes were not statistically enriched in Germline_Lv3 when compared with other germline populations, expression levels of germline-enriched genes such as *Lv-Nan2, Lv-Ttk1* and *Lv-Cecr2L* were comparable with that of Germline_Lv1 (Fig. 3C and S3). These results strongly suggest that Germline_Lv1 and Germline_Lv3 were *bona fide* germline cell populations. Upregulated ribosomal gene expression was observed in Germline_Lv3 (Fig. 3C). Indeed, the top5 marker genes include Lv-Rpl22, Lv-Rpl11, Lv-Rps15a, Lv-Rpl28 and Lv-Rps15 (Table S7). Further, the top-50 marker genes include 39 ribosomal genes (Table S7). However, since the upregulation of ribosomal genes was observed throughout the embryo (Fig. S4), the upregulation is a general phenomenon observed throughout the *Lv* embryo rather than a germline-specific phenomenon. In addition, endodermal gene expression was also upregulated in the Germline_Lv3 compared to other germline cell populations (Fig. 3C and S5). These results suggest that the small PGC population in these embryos has diverse biology that may influence differentially their future fate.

### Integration of both scRNA-seq datasets from *Sp* and *Lv* embryos

We employed the batch-effect correction to enable a direct comparison between this *Lv* dataset and the *Sp* dataset (Foster *et al*., 2020). We integrated the *Lv* dataset with the *Sp* dataset using four developmental time points individually (EB–LG; *Lv-Sp* dataset) (Fig.S6A–D) and then normalized by Liger, another batch-effect correction tool (Fig. S6I–L) (Welch, 2018). Harmony did not correct the batch effects between *Lv* and *Sp* datasets effectively (Fig. S6E–H)

In the *Lv-Sp* datasets, we identified seven to nine ectodermal cell populations (Clusters EB2–EB8, MB2–9, EG2–8 and LG2–10; Fig.4A–H). In addition, we showed the expression patterns of *Ebr1, Ac/Sc* and *Brn1-2-4* as ectodermal marker genes (Fig. 4B, D, F and H). *Ebr1* was enriched in dorsal ectoderm in the *Lv* dataset (Cluster Lv7–9) (Fig.1C, Table S2). *Ac/Sc* and *Brn1-2-4* are known to be expressed in proneural cells during embryogenesis (Slota and McClay, 2018). One to two endodermal cell populations were identified by enriched expression of *FoxA, Blimp1, Endo16, Gatae, Bra-1, Hox11/13b* and *Eve* (Fig.4B, D, F and H). Consistent with the results of the *Lv* dataset, *Lv*-derived endodermal cells in EB and MB (EB9 and MB11) showed weaker *Endo16* expression than that in EG and LG embryos (Fig. 4B, D, F and H). In *Sp* embryos, the *Endo16* expression was still low until EG stage (EG9 and 10) (Fig. 4F). Further, an endodermal cell population lacking *Blimp1* expression was observed in both *Lv* and *Sp* in LG embryos (LG12) like the Lv18 in the *Lv* dataset. Three mesodermal cell populations in EB, MB and LG embryos and two mesodermal cell populations in EG embryos were identified by significantly enriched mesodermal gene expression (Fig. 4B, D, F and H; Table S3–S6). Furthermore, endodermal genes, *Blimp1, Endo16* and *Gatae*, were enriched in secondary mesenchyme cell (SMC) populations in EB stage (EB12) (Fig. 4B; Table S3). Since EB12 did not show enriched *FoxA* expression (Fig. 4A and B; Table S3), we identified EB12 as a presumptive SMC population.

**Fig.4.**
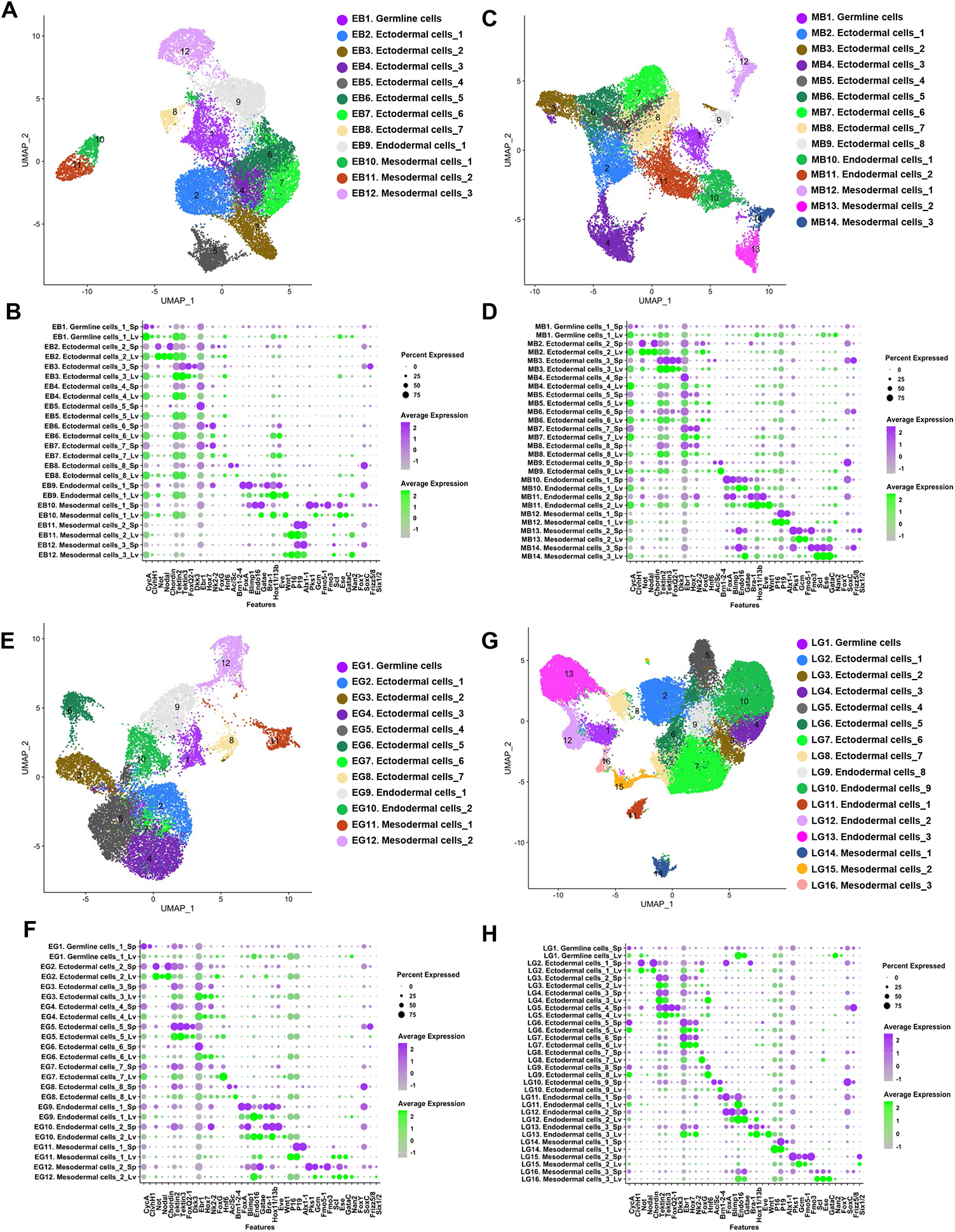
Marker gene expression in the *Lv-Sp* datasets. (A–H) UMAP plots (A, C, E and G) and dot plots (B, D, F and H) for EB (A and B), MB (C and D), EG (E and F) and LG (G and H) embryos are presented. Dot plot represents marker gene expression in *Lv*-derived cells (green) and *Sp*-derived cells (purple) separately. Dot size and dot color indicate the percentage of cells expressing the gene and the average expression level, respectively.

### PGC comparison between *Lv* and *Sp*

In the *Lv-Sp* dataset, *Nanos2* expression was observed not only in the germ cells but also in somatic cells of the *Lv* embryo (Fig. S7). Further, considering that the PGCs represent a relatively rare population compared to somatic cells, too many cells were classified as the germ cell clusters (EB1, MB1, EG1 and LG1). These observations suggest that the germ cell clusters likely contain non-germline cells. In the Lv-dataset, Nanos2-FoxY double positive cluster (Lv18; Veg2 mesoderm) and Nanos2 positive cluster (Lv23; PGCs) were identified in LG embryos, but only Nanos2-FoxY double positive cluster (LG1) was found in the LG embryo of the Lv-Sp dataset, suggesting that LG1 includes both PGCs and Veg2 mesoderm and these two populations were not distinguishable in *Lv-Sp* dataset probably due to the similar marker gene expression. FoxY is an essential marker in this analysis since it was found to be an essential transcription factor to make *Sp* Nanos2 in the veg2 cells. Therefore, *Nanos2*-positive cells were extracted from the *Lv-Sp* datasets and reanalyzed in detail (Fig.5). These *Nanos2*-positive cells were grouped into seven subpopulations by UMAP plot (Nan2pos_1–7; Fig. 5A-C) and were further separated into two datasets containing either *Lv*-derived (Fig. 5B) or *Sp-*derived (Fig. 5C) cells.

**Fig.5.**
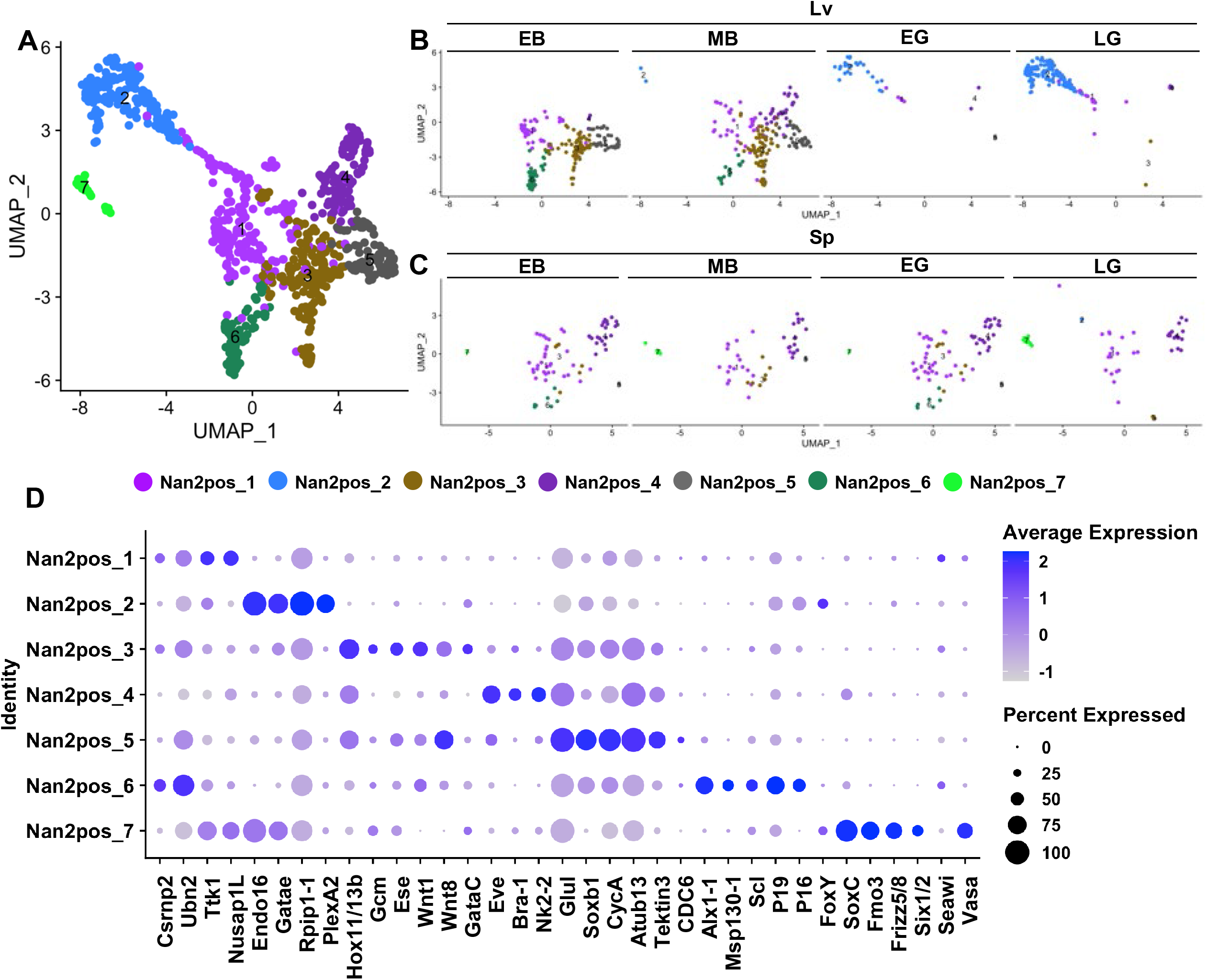
Germline subpopulations in the *Lv-Sp* dataset. (A–C) UMAP plots for Nanos2 positive cells of the *Lv-Sp* datasets. UMAP plots of *Lv*-derived cells (B) and *Sp*-derived cells (C) in each developmental stage are presented. (D) Dot plot showing marker gene expression in Nanos2-positive cells.

Among the seven subpopulations, Nan2pos_1 is the most likely to be the PGCs for the following reasons: (1) Nan2pos_1 showed enriched expression of *Csrnp2, Ttk1* and *Nusap1L*, all of which were expressed in the Germline_Lv1 of the *Lv* dataset, (Donoughe *et al*.) Nan2pos_1 was detected throughout embryogenesis (Fig. S9A–B). Nan2pos_5 and Germline_Lv2 were expressing common marker genes, *Soxb1, CycA, Atub13* and *CDC6* (Fig. 3C and 5F; Table S7 and S8). These results show that Nan2pos_1 and Nan2pos_5 correspond to Germline_Lv1 and Germline_Lv2, respectively. In the *Lv* dataset, Germline_Lv3 was found as the germ cell population in EG and LG embryos (Fig. 3A–C) and Nan2pos_2 shows similar marker gene expression with Germline_Lv3. Endo16, Gatae, Rpip1-1 and PlexA2 are enriched both in Germline_Lv3 and Nan2pos_2. However, in contrast to Germline_Lv3, Nan2pos_2 expresses FoxY, suggesting that the Nan2pos_2 cluster is either the Veg2 mesoderm or a mixture of Veg2 mesoderm and PGCs.

In addition, somatic cells were identified based on well-studied marker gene expression as follows (Fig. 5D): Nan2pos_3 (SMCs), Nan2pos_4 (oral ectoderm), Nan2pos_6 (PMCs) and Nan2pos_7 (Veg2 mesoderm). These somatic cell populations were observed not only in *Lv* but also in *Sp* embryos (Fig. 5C and D). These results suggest that, although *Nanos2* was predominantly expressed in the PGCs in *Sp* embryos (Fig. S7) (Foster *et al*., 2020), *Nanos2* is also slightly but significantly expressed in somatic cells. Furthermore, Nan2pos_7 (veg2 mesoderm) expressing *FoxY, SoxC, Fmo3* and *Frizz5/8* cells were observed only in *Sp* embryos (Fig. 5D).

### NPDE-mediated post-transcriptional regulation of *Nanos2*

Considering that *Nanos2* mRNA was widely expressed in *Lv* embryos compared to *Sp* embryos (Fig. 2A–H and Fig. S7A–H), *Lv Nanos2* is most likely to be regulated translationally and/or post-translationally to establish agermline-specific Nanos2 protein expression. In *Sp* embryos, *Nanos2* expression is post-translationally regulated by the Nanos Protein Degradation Element (NPDE) (Oulhen and Wessel, 2016). Therefore, we aimed to investigate whether the NPDE sequence is conserved among sea urchin species. For this purpose, *Nanos2* protein sequences were compared between *Lv, Sp* and *Hemicentrotus pulcherrimus* (Colonnetta *et al*.), a sea urchin species closely related to *Sp* (Fig. S10). Although Nanos2 is highly divergent in sequence between animals, the NPDE sequence is highly conserved between *Sp* Nanos2 and *Hp* Nanos2 (Fig. S10). Although the *Lv* Nanos2 protein contains indels in the NPDE, the C-terminal region is completely conserved (Fig. S10, aa 36–46; ITELSKVMRG). These results led us to hypothesize that the well-conserved C-terminal region is functionally important among these species.

To test this hypothesis, we divided the *Sp* Nanos2 NPDE element into eight fragments consisting of 13–22 amino acids (Fig. 6A) and the corresponding nucleotide sequences were fused to the GFP ORF and *Sp Nanos2* 3’UTR without the GNARLE sequence (*ΔGNARLE*). The GNARLE is required to restrict mRNAs to the PGCs (Oulhen *et al*., 2013). The mRNAs containing *Sp Nanos2* 3’ UTR *ΔGNARLE* are not regulated post-transcriptionally, and in turn are distributed in both somatic and germ cells. These *in vitro* synthesized mRNAs were injected into *Sp* embryos, together with *mCherry* mRNA. The expression of these fused proteins (GFP-NPDEFL and GFP-NPDE1–8) were observed in blastula stage embryos and then normalized by the signal intensity of mCherry protein. The normalized signal intensity of GFP-NPDE fusion proteins were compared with control GFP (Fig. 6B, B’ and L). As reported previously, expression of the full length NPDE (GFP-NPDEFL) was barely detectable (Fig. 6C and C’) (Oulhen and Wessel, 2016). In addition, fusion with NPDE4–8 significantly decreased the GFP intensity compared to the control GFP, while fusion with NPDE1–3 did not attenuate GFP intensity (Fig. 6D–L and 6D’–K’). Notably, fragments including the highly conserved C-terminal region (GFP-NPDE5 and 8) exhibited a significant decrease in GFP intensity but were never comparable to the full length NPDE (Fig. 6C, C’, K and K’). These results suggest that the C terminal region of the NPDE (Fig. S10; amino acids 36 to 45) is important but not sufficient to regulate Nanos2 protein expression.

**Fig.6.**
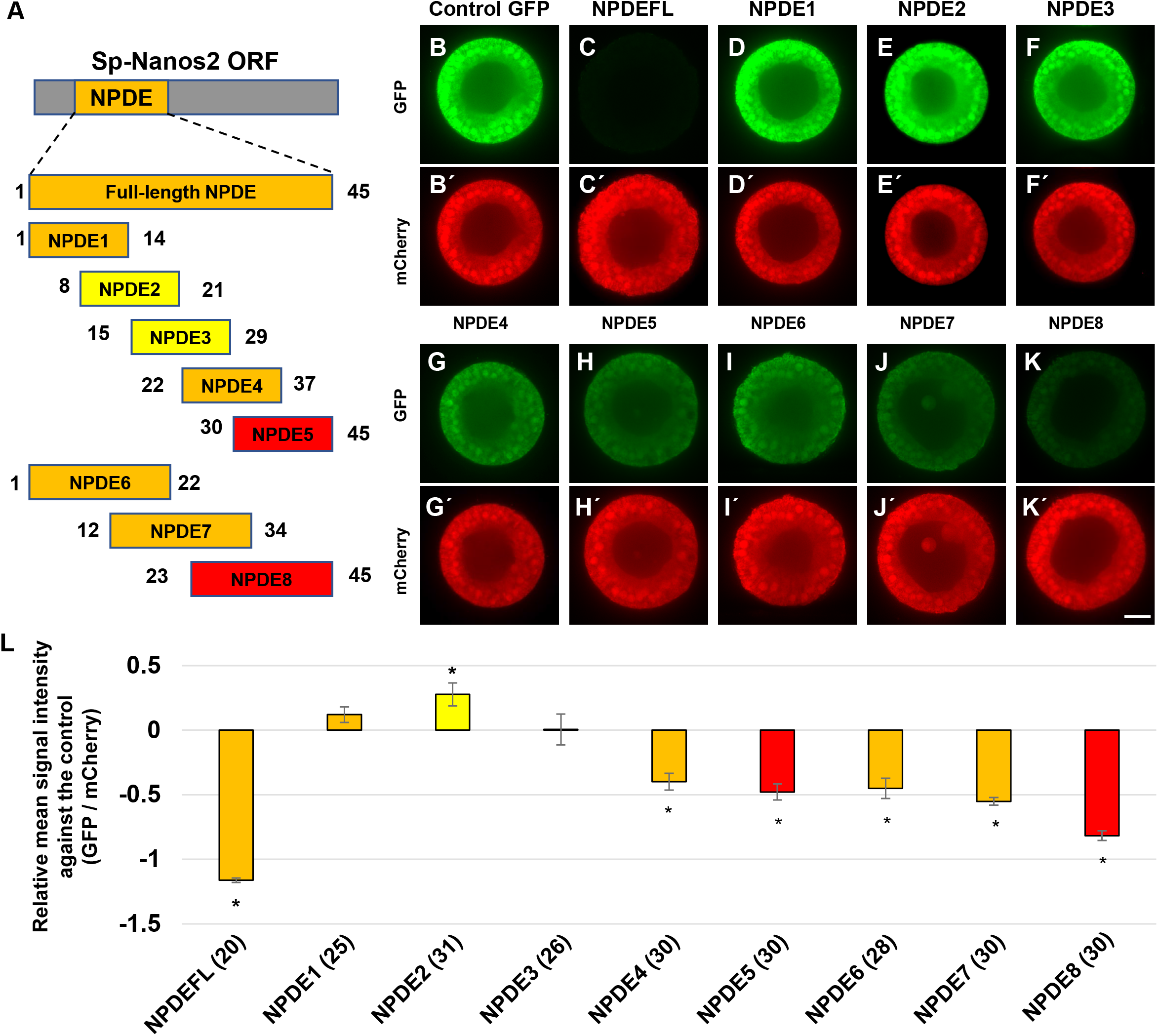
NPDE-mediated protein degradation in Sp embryos. (A) Full-length NPDE (orange box) located at the N-terminal side of *Sp Nanos2* ORF (gray box) was divided into eight fragments (NPDE1–8). Amino acid position within the NPDE is shown at both side of each fragment. The fragments are color-coded by the percentage of amino acid residue conserved in *Lv Nanos2* (red: >60%; orange: 30∼60%; yellow: <30%). (B–K and B’–K’) *Sp* embryos injected with GFP and mCherry mRNAs under the control of β-globin UTRs. Fluorescence of GFP (B–K; green) and mCherry (B’–K’; red) were observed in the EB embryos. Scale bar: 20 µm. (L) Relative mean signal intensity against the control is presented. Graph is color-coded as described above. Significance was calculated between control GFP and GFP-NPDE fusion proteins (NPDEFL and NPDE1–8) by Student’s t test (*: p < 0.05). Error bars indicate standard errors of samples. The numbers of injected embryos examined are shown in parentheses.

### Regulation of *Lv Nanos2* expression

To determine whether the cis- and trans-regulatory mechanisms of *Sp Nanos2* mRNA are conserved in *Lv*, we injected mRNAs encoding *Sp Nanos2* ORF and *Sp Nanos2* 3’ UTR (*Sp-Sp*), *Lv Nanos2* ORF and *Lv Nanos2* 3’ UTR (*Lv-Lv*), and the hybrid mRNAs (*Sp-Lv* and *Lv-Sp*) into *Lv* embryos. GFP ORF fused with β-globin UTRs (control GFP) was used as the control. We observed the GFP mRNA and protein expression at blastula stage (Fig. 7). Control-GFP, *Lv-Sp* and *Sp-Sp* did not show enriched GFP RNA or protein in the *Lv* embryos (Fig. 7A–A’’, C–C’’ and E–E’’). These results suggest that the regulatory elements of *Sp Nanos2* are not functional in *Lv* embryos. In addition, and as expected by the low overall conservation of the NPDE of *Lv* Nanos2, we discovered that the *Lv Nanos2* ORF is not sufficient for its protein enrichment in the PGCs (Lv-Sp). Interestingly, when mRNAs containing *Lv Nanos2* 3’UTR (*Lv-Lv* and *Sp-Lv*) were injected into these *Lv* embryos, the translated protein was enriched in the PGCs, while the corresponding RNAs were not (Fig.7B–B’’ and D–D’’). This result shows that the *Lv Nanos2* 3’ UTR is sufficient to establish germline-specific expression by translational regulation in the *Lv* embryos. Further, when the *Lv-Lv* mRNA was injected into *Sp* embryos, the expression pattern of the injected mRNA and resulting protein were completely the same with that of control GFP (data not shown). These results show that *Lv* and *Sp* employ different regulation mechanisms and both the cis- and trans-regulatory elements for Nanos2 were not conserved.

**Fig.7.**
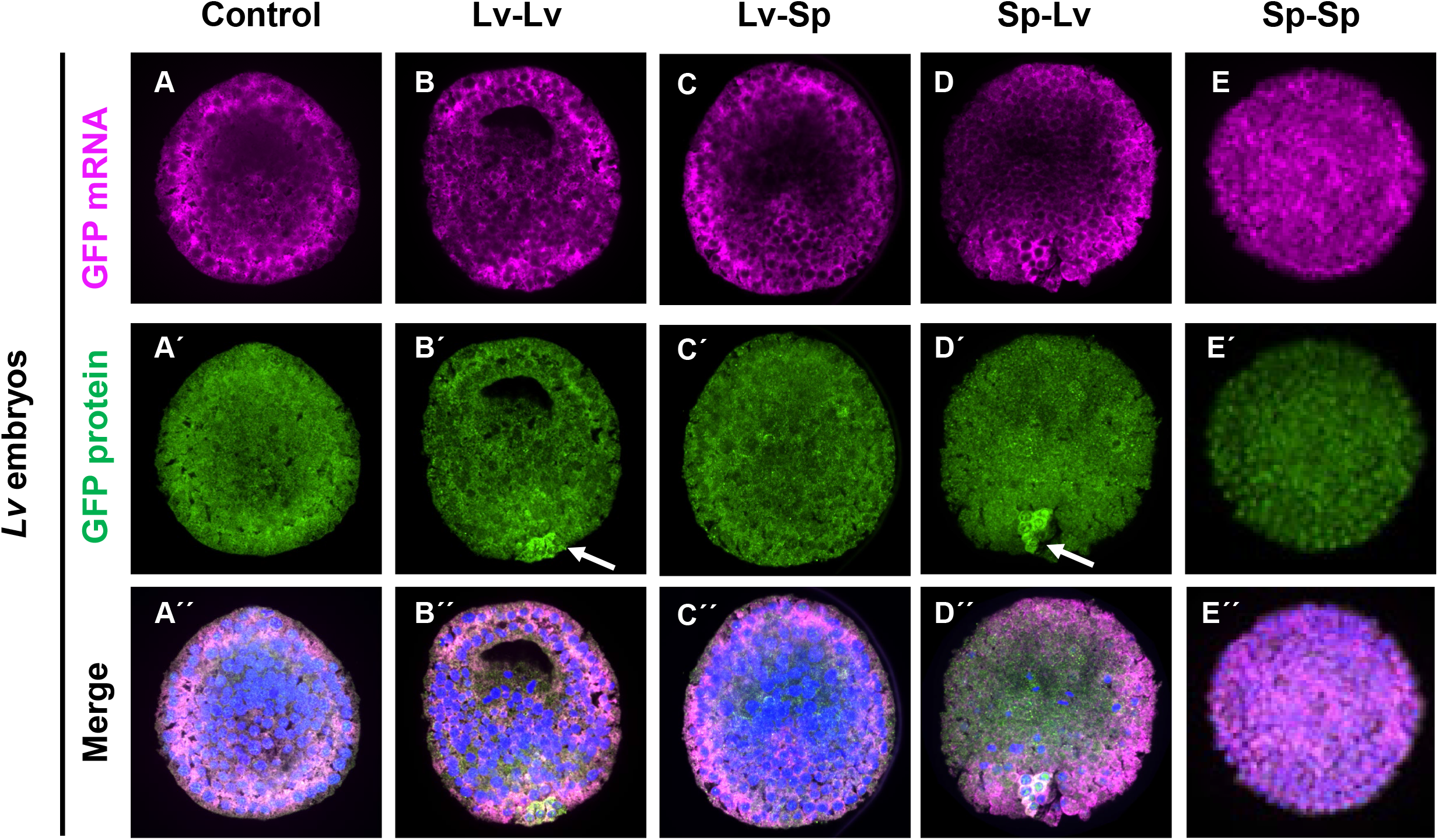
*Lv* Nanos2 3’UTR regulates its protein expression. *Lv* embryos injected with Control GFP mRNA (A), *Lv-Lv* mRNA (B), Lv-Sp mRNA (C), Sp-Lv mRNA (D) and Sp-Sp mRNA (E) were labelled by *in situ* hybridization for GFP mRNA (magenta; A–E) and immunohistochemistry for GFP protein (green; A’–E’). DNA staining (blue) merged with in situ hybridization and immunostaining are presented (A’’–E’’). White arrows in B’ and D’ indicate the specific translation of the constructs in the germ cells. Scale bar: 20 µm.

## Discussion

### Germline gene expression in *Lv* embryos

We found that the transcriptional profile of the PGCs in Lv was quite similar with that of undifferentiated cells of the early embryo, and they contained no prominent marker genes of germ cells except for the *Lv* Nanos2. This may be explained by previous studies in the closely related sea urchin species, *Strongylocentrotus purpuratus*. In *Sp* CNOT6 transcripts encoding a deadenylase is depleted from the PGCs which creates a stable environment for maternally inherited mRNAs (Swartz *et al*., 2014). This stable environment is likely to be established in both *Lv* and *Sp* germ cells since maternal mRNAs are also retained in *Lv* germ cells. In *Sp*, transcriptional, RNA degradation and translational activities are globally repressed in the PGCs (Swartz *et al*., 2014, Oulhen *et al*., 2017). Quiescence is a widely conserved feature of germ cells to protect them from somatic differentiation by repressing somatic gene expression. Considering that *Sp* Nanos2 is the key regulator to establish quiescence, *Lv* Nanos2 may be responsible for the transcriptional profile of the germ cells.

Our results show the presence of two germline subclusters throughout the development, an early Germline_Lv1 and a later Germline_Lv3. In the *Sp* sea urchin, four PGCs are formed early during the development, their cell cycle is downregulated until gastrulation when the level of *Sp* Nanos2 starts to decrease and that the PGCs finally divide once to give rise to 8 cells. These 2 subclusters could represent the quiescent PGCs (Germline_Lv1), becoming more active during gastrulation (Germline_Lv3). Comparing the differential gene expression between these two subclusters overtime could reveal new mechanisms of how PGCs exit quiescence during gastrulation.

### Comparison of Nanos2 regulation between *Sp* and *Lv* sea urchins

In this study, we found that *Lv Nanos2* RNA is expressed not only in the PGCs but also in the somatic cells including three germ layers. In the sea urchin *Sp*, Nanos2 mRNA and protein are highly restricted to the PGCs at blastula stage. *Sp* Nanos2 is transcribed broadly by the Wnt pathway (Pieplow *et al*., 2021), but the resulting mRNA is quickly degraded outside of the PGCs and retained only in the PGCs. The presence of an element, GNARLE, on its 3’UTR is required for this regulation (Oulhen *et al*., 2013). In contrast, *Lv* Nanos2 mRNA is not restricted to the PGCs and its 3’UTR does not have a conserved GNARLE. In *Sp*, Nanos2 is also regulated at the protein level by the NPDE (Nanos protein degradation element). This element is not highly conserved and is not functional in *Lv* embryos (Fig S11). We discovered that instead of relying on RNA and/or protein stability, the germline expression of *Lv* Nanos2 mostly depends on translational regulation. The presence of the *Lv Nanos2* 3’UTR on an exogenous RNA leads to a significant restriction of the corresponding protein in the PGCs. We conclude that whereas its mRNA is present broadly in the embryos, it is predominantly translated in the PGCs. Nanos has previously been shown in other animals to also be regulated at the translational level: in *Xenopus, C. elegans*, and *Drosophila* (Gavis and Lehmann, 1992, Subramaniam and Seydoux, 1999, Forrest and Gavis, 2003, Forrest *et al*., 2004, D’Agostino *et al*., 2006, Kalifa *et al*., 2006, Jadhav *et al*., 2008, Luo *et al*., 2011). Taken together, translational regulation is most likely to be the basal mechanism widely conserved between animals including *Lv*. By contrast, post-transcriptional and post-translational regulations play a key role in *Sp*. The selection for these diverse regulatory mechanisms is still unknown.

### Somatic cell gene expression

*Nanos2* expression was not the only difference observed between both datasets. Several somatic genes were also differentially expressed between *Sp* and *Lv*. For example, endodermal cell populations were identified by enriched expression of *FoxA, Blimp1, Endo16, Gatae, Bra-1, Hox11/13b* and *Eve* in the *Lv-Sp* dataset (Fig. 4). At the early gastrula stage, Endo16 is more abundant in *Lv* compared to *Sp* (cluster EG9). Similar results were observed in the ectodermal clusters. An obvious example is Hnf6 (cluster EG7), that is also found more abundantly in *Lv* compared to *Sp*. These are just a few examples suggesting that different mechanisms could be also used by these two sea urchins to regulate the expression of their somatic genes through development.

### Nanos2 mRNA was detected in the Veg2 mesoderm

In the *Lv-Sp* dataset, seven subpopulations of Nanos positive cells have been characterized (Table S8). Importantly, Nan2pos_7 is only detected in the *Sp* sea urchin. Nan2pos_2 is only detected in the *Lv* sea urchin. Both of these populations reached their peak of abundance at the late gastrula stage. They both express markers such as *FoxY* and *Endo16*, suggesting that both of these populations could represent the Veg2 mesoderm. In *Sp* embryos of which micromeres, parent cells of PGCs, are surgically removed, new germline cells are regenerated during embryogenesis and then most of the resulting adults generate functional gametes (Yajima and Wessel, 2011). Our previous data also suggested that in *Sp*, the Veg2 mesoderm could be important to form a new germline after micromere removal (Voronina *et al*., 2008, Oulhen *et al*., 2019b). Micromere-deleted *Lv* embryos can also regenerate their germline but use distinct mechanisms (Voronina *et al*., 2008). For example, in contrast to *Lv*, micromere-deleted *Sp* embryo shows significant upregulation of Vasa protein expression. Our data here suggest that these two sea urchins could respond differently to micromere removal because of their differential gene expression in the Veg2 mesoderm that do not overlap in the umap: *Sp* Nanos2pos_7 and *Lv* Nanos2pos_2 (Fig 5). Nanos2 expression may be optimized for germline regeneration in *Strongylocentrotus purpuratus* embryos.

### Evolution of Germline gene regulation

We showed that *Nanos2* is differentially expressed between both urchins (Fig 8). It is specific to the germline cluster in *Sp*, but it is more broadly expressed in *Lv*. However, this observation is not a common characteristic of all the germline genes. For example, the expression of Vasa and Seawi transcripts do not follow the same pattern; instead, their expression is similar in both sea urchins (Fig S12). Nanos ectopic expression in embryos is toxic (Luo *et al*., 2011), so animals have found various methods to restrict its expression to the germ cells. In both sea urchins, the 3’UTR of Nanos2 is essential but the regulations are different. In *Sp*, this 3’UTR is essential to degrade Nanos2 mRNA in the somatic cells, but in *Lv*, this 3’UTR instead leads to its specific translation in the germ cells. How these regulations happen is still unknown. Further studies will include the identification of the proteins associated with each of these 3’UTRs.

**Fig.8.**
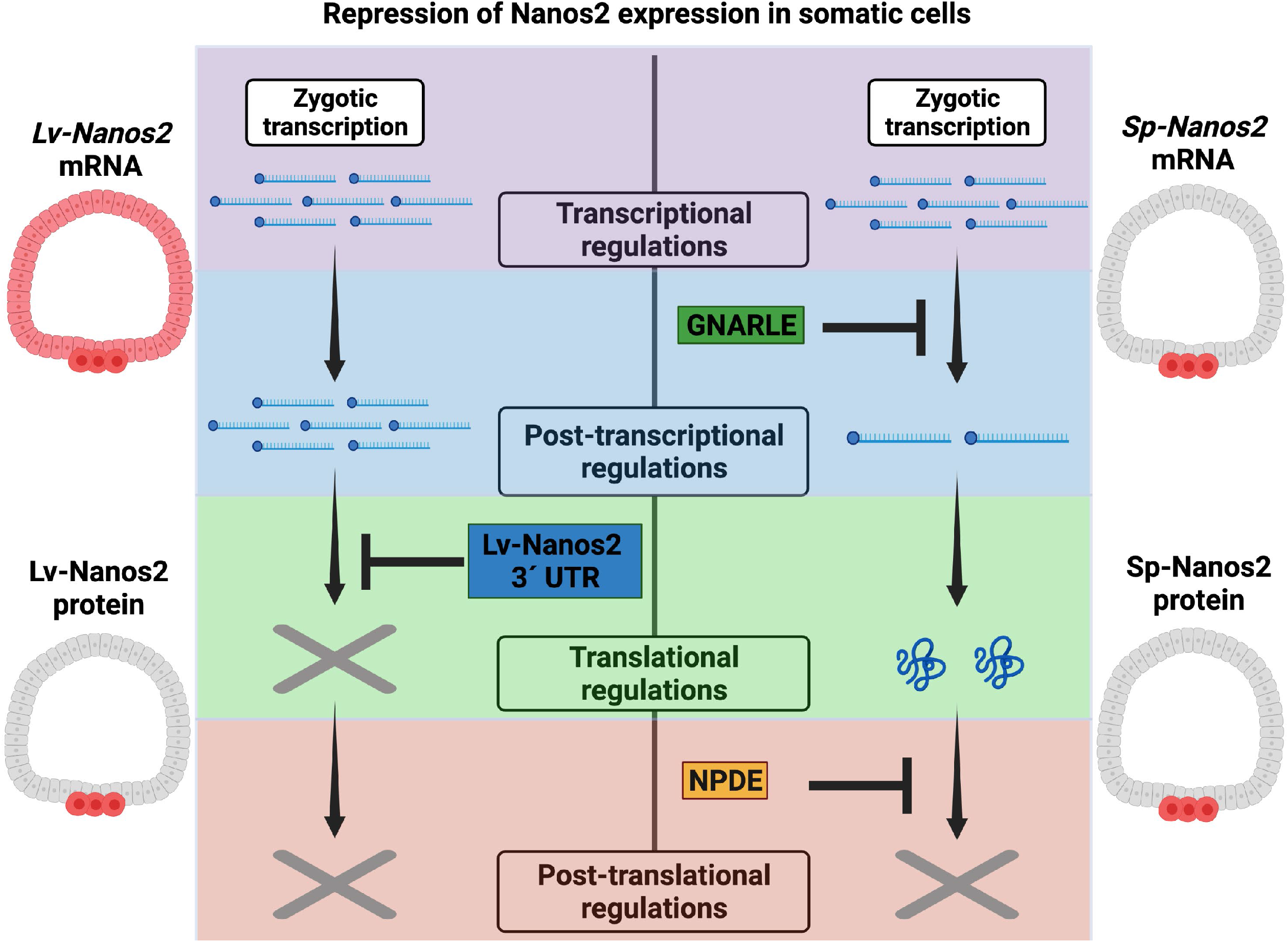
Distinct regulations of Nanos2 expression in sea urchin. In *Lv*, Nanos2 mRNA is broadly expressed. Its 3’UTR leads to a specific protein synthesis in the germline. In *Sp*, Nanos2 mRNA is degraded in the somatic cells through an element in its 3’UTR (GNARLE). Sp Nanos2 protein stability is regulated through the NPDE, Nanos protein degradation element, and leads to its degradation in the somatic cells and its retention in the germ cells.

## Materials and Methods

### Animal care and culturing

Adult *Strongylocentrotus purpuratus* were obtained from Pete Halmay of Pt. Loma Marine Invertebrate Lab (Lakeside, CA) and housed in aquaria with artificial seawater (ASW) at 16°C (Coral Life Scientific Grade Marine Salt; Carson, CA). Adult *Lytechinus variegatus* were obtained from the Pelagic Corporation and from the Duke University Marine Laboratory, Beaufort NC. They were housed in aquaria with artificial seawater (ASW) at room temperature (Coral Life Scientific Grade Marine Salt; Carson, CA). Eggs and sperm of *S*.*purpuratus* or *L*.*variegatus* were spawned by injection of 0.5M KCl into the adult coelomic cavity. Fertilization was accomplished in sea water containing 1mM 3-Amino-1,2,4-triazole to reduce cross-linking of the fertilization envelope, and which was washed out after 30 minutes. Embryos were cultured in filtered (0.2micron) sea water collected at the Marine Biological laboratories in Woods Hole MA, until the appropriate stage. *Lv* embryos were cultured at room temperature, *Sp* embryos were cultured at 15°C.

### *Lv* culture and dissociation for scRNA-seq

All embryos used in the study to obtain the *Lv* scRNA-seq dataset resulted from mating of one male and one female. Multiple fertilizations were initiated in this study and timed such that the appropriate stages of embryonic development were reached at a common endpoint. The embryos were then collected and washed twice with calcium-free sea water, and then suspended hyalin-extraction media (HEM) for 10-15 minutes, depending on the stage of dissociation. When cells were beginning to dissociate, the embryos were collected and washed in 0.5M NaCl, gently sheared with a pipette, run through a 40micron Nitex mesh, counted on a hemocytometer, and diluted to reach the appropriate concentration for the scRNA-seq protocol. Equal numbers of embryos were used in each time point and at no time were cells or embryos pelleted in a centrifuge (Oulhen *et al*., 2019a).

### Genome-guided de novo transcriptome assembly

For de novo transcriptome assembly, whole embryo RNA-seq dataset (Li *et al*., 2020) were downloaded from NCBI (https://www.ncbi.nlm.nih.gov): SRR9673381, SRR9673382, SRR9673383, SRR9673384, SRR9673385, SRR9673386, SRR9673387, SRR9673388, SRR9673403, SRR9673404, SRR9673405, SRR9673406, SRR9673407, SRR9673408, SRR9673409 and SRR9673410. The raw reads were preprocessed by Trimmomatic and then mapped to the *Lv* genome sequence from Li et al, (2020) by using Hisat2. The output SAM files were converted to BAM files and merged into a single BAM file. To obtain transcriptome data, the BAM file was assembled *de novo* using Trinity with default settings.

### Prediction of UTR information

We used a gene annotation file from Li et al, (2020) with some modification. Since the annotation file contains only CDS regions, the UTR regions were predicted by Program to Assemble Spliced Alignments (PASA) based on the transcriptome information described in “Genome-guided de novo transcriptome assembly”. In order to maximize the incorporation of UTR information into gene annotation, we performed PASA-pipeline two times. The PASA-updated annotation GFF3 file was converted into GTF file format by gffread.

### Prediction of conventional gene names

To identify the conventional gene names of contigs in PASA-updated annotation files, we built a BLASTP database with a *Sp* peptide sequences downloaded from echinobase (S. purpuratus v5.0). The Lv peptide sequences from Li et al, (2020) was used for the query sequence for BLASTP analyses and best hit genes were identified as SPU_IDs. The SPU-IDs were converted into conventional gene names by using a table from echinobase (GenePageGeneralInfo_AllGenes.txt). The gene names in PASA-updated annotation file were replaced with the identified conventional gene names.

### Single cell RNA sequencing

Single cell encapsulation was performed using the Chromium Single Cell Chip B kit on the 10x Genomics Chromium Controller. Single cell cDNA and libraries were prepared using the Chromium Single Cell 3’ Reagent kit v3 Chemistry. Libraries were sequenced by Genewiz on the Illumina Hiseq (2×150 bp paired-end runs). Single cell unique molecular identifier (Fujii *et al*.) counting was performed using Cell Ranger Single Cell Software Suite 3.0.2 from 10X Genomics. Duplicate blastula and gastrula stage libraries were aggregated using the cellranger aggr function. Cellranger gene expression matrices were further analyzed using the R package Seurat v 3.1.4 (Butler *et al*., 2018, Stuart *et al*., 2019). A Seurat object was created by simply combining the individual datasets and then normalized by scaling gene expression in each cell by total gene expression. The top 2000 highly variable genes were then used for downstream analysis. The batch effects between samples were corrected by Harmony (Korsunsky *et al*., 2019) with default setting. UMAP analysis was performed with the following parameters: dims = 1:20, resolution = 1.

### Comparison between *Lv* and *Sp* datasets

To compare transcriptional profiles between *Lv* and *Sp*, 993 genes from the *Lv* dataset and 1218 genes from the *Sp* dataset were selected with adjusted p value < 1.00E-200. In addition, about 1000 genes that may play important roles during development were added manually. Next, the homologous genes between *Lv* and *Sp* were identified by checking the reciprocal best hits of BLASTP analysis. Genes without reciprocal best hit were not used for the downstream analysis because the genes are likely to be species-specific and not suitable for inter-species comparison. Finally, 886 genes were identified as the homologous genes between *Lv* and *Sp*. The homologous gene information was extracted from *Lv* annotation file (Li et al., 2020) and *Sp* annotation file (*S. purpuratus* v5.0) and then the gene names were unified with exactly the same conventional gene names in both annotation files. Sequenced reads from *Lv* and *Sp* embryos were mapped to corresponding genome sequence and then counted based on the newly made annotation files containing 886 homologous genes. The resulting datasets were combined between species and then normalized by scaling gene expression in each cell by total gene expression. The batch effects between species were corrected by Liger with the following settings: k = 20, lambda = 5. UMAP analysis was performed with the following parameters: dims = 1:20, resolution = 0.5. These interspecies comparisons were performed using EB, MB, EG and LG embryos separately.

### Plasmid construction

For construction of GFP-NPDEFL and GFP-NPDE1–8 plasmids, *Sp* Nanos2 5’ UTR, *Sp* Nanos2 3’ UTR ΔGNARLE, GFP ORF and NPDE fragments were amplified from Nanos2-GFP ΔGNARLE (Oulhen and Wessel, 2016). These amplicons were cloned into pGEM-T Easy Vector using NEBuilder. For construction of plasmids containing *Sp Nanos2* ORF and *Sp Nanos2* 3’ UTR (*Sp-Sp*), *Lv Nanos2* ORF and *Lv Nanos2* 3’ UTR (*Lv-Lv*), and the hybrid (*Sp-Lv* and *Lv-Sp*), total RNAs were isolated from *Lv* and *Sp* embryos using TRIzol, and cDNAs were synthesized using SuperScript III Reverse Transcriptase (Thermo Fisher Scientific, Cat#18080093). The ORF and 3’ UTR of *Lv* Nanos2 and 3’ UTR of *Sp* Nanos2 were amplified from the cDNAs. ORF and 5’ UTR of *Sp* Nanos2 and GFP ORF were amplified as described above.

### Microinjections

Sea urchin eggs were dejellied by washing for 10 minutes in pH5.0 seawater (*Sp*), or by passing the eggs through a 100uM mesh (*Lv*). Eggs were then rowed on protamine sulfate coated Petri dishes. Zygotes were injected with 2 pL of injection solution, by constant pressure and in the presence of 1 mM 3-AT (Sigma). The injected zygotes were cultured at 16C for *Sp*, or at Room Temperature for *Lv*.

## Data availability

The sequencing files and gene expression matrices for the single-cell RNA-seq analysis presented here have been deposited in the GEO database and may be accessed GSE208709.

## Acknowledgments

We are grateful to the NIH (1R35GM140897, GMW and 1P20GM119943, NO), the NSF (IOS-1923445, GMW) and the TOYOBO Biotechnology Foundation long term research grant (to SM) for the support to conduct this research.

## Table legends

**Table.S1 The number of cells analyzed in the Lv-dataset**

Each developmental stages and the total cell number is shown separately.

**Table.S2 Marker genes identified in the Lv-dataset**

**Table.S3 Marker genes identified in the EB embryos of Lv-Sp dataset**

**Table.S4 Marker genes identified in the MB embryos of Lv-Sp dataset**

**Table.S5 Marker genes identified in the EG embryos of Lv-Sp dataset**

**Table.S6 Marker genes identified in the LG embryos of Lv-Sp dataset**

**Table.S7 Marker genes identified by sub-clustering of the germline cells of the Lv dataset**

**Table.S8 Marker genes identified by sub-clustering of the Nanos2-positive cells of the Lv-Sp dataset**

**Fig.S1.**
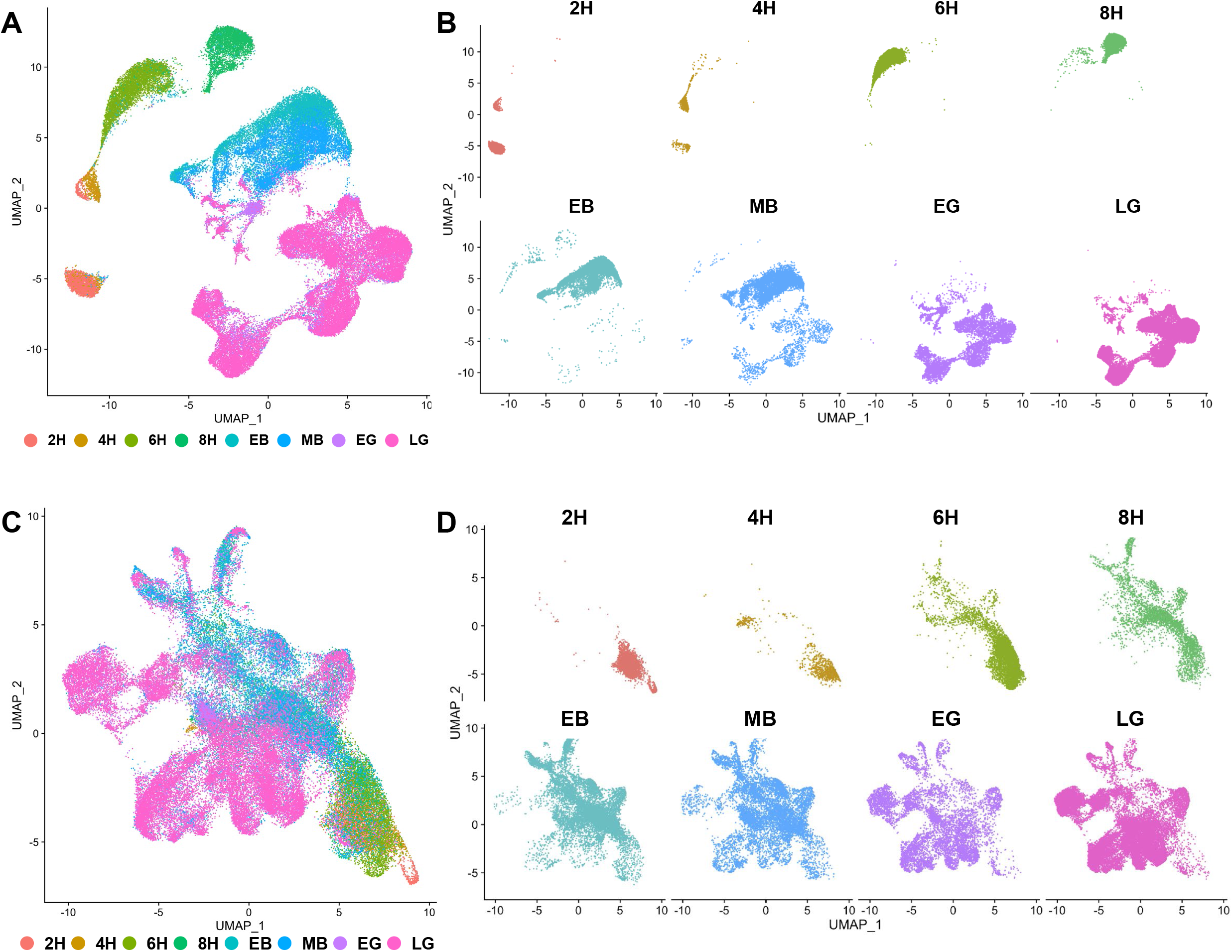
UMAP plot analysis of non-normalized and Harmony-normalized *Lv-*dataset. (A–D) Non-normalized *Lv*-dataset (A and B) and Harmony-normalized *Lv*-dataset (C and D) are presented. Cells are color-coded by the developmental stage (2H–LG). Datasets are split into each developmental stage (B and D).

**Fig.S2.**
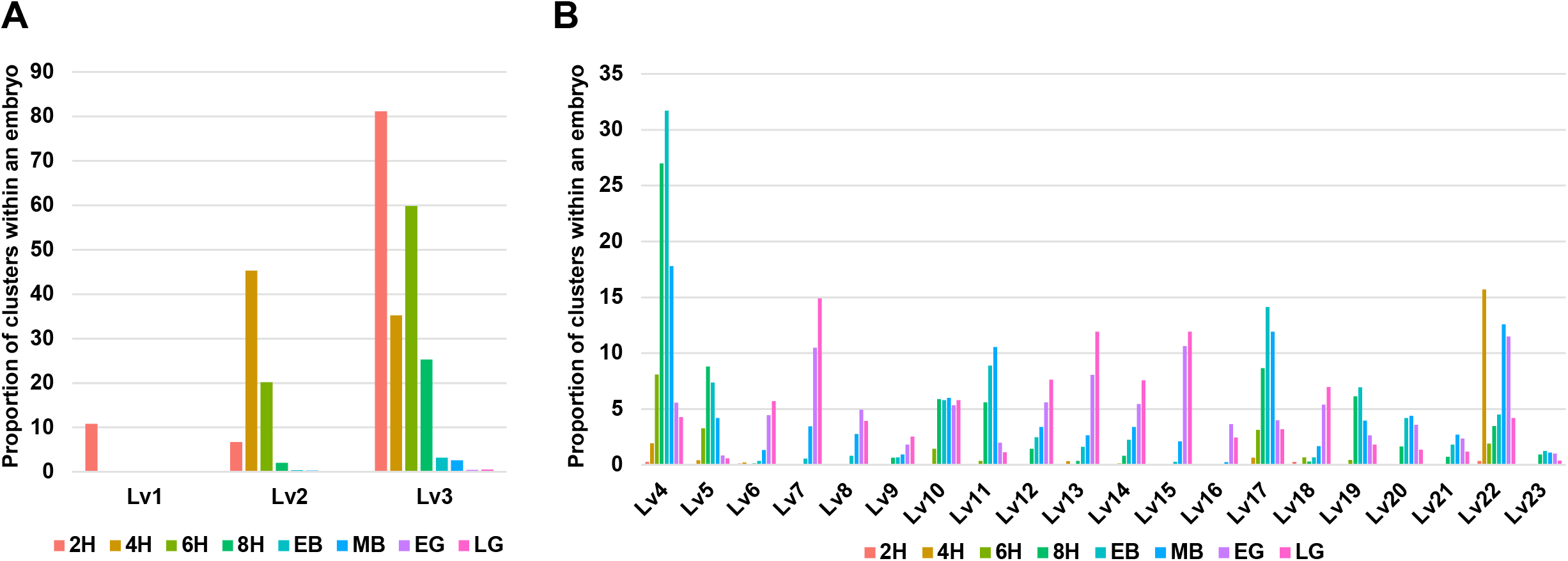
Identification of cell populations in *Lytechinus variegatus* embryos. The proportion of undifferentiated cell populations (A) and ectoderm, endoderm mesoderm and germ cells (B) within an embryo are shown. The proportions were calculated by using the total number of cells analyzed by scRNA-seq for each developmental stage.

**Fig.S3.**
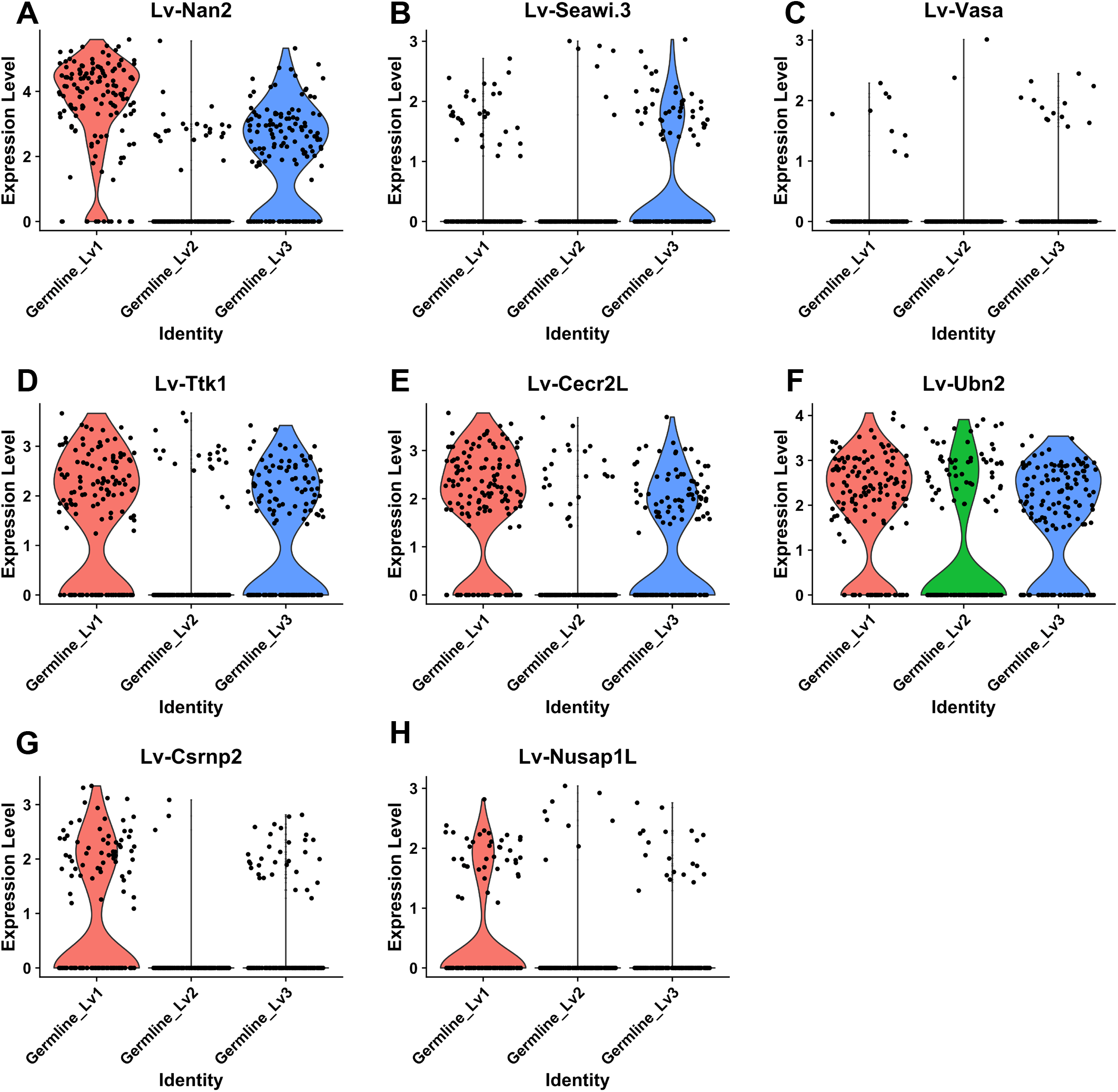
Marker gene expression in the germline subpopulations of *Lv* dataset. Violin plots showing marker gene expression in Germline_Lv1 (Chatfield *et al*.), Germline_Lv2 (green) and Germline_Lv3 (blue).

**Fig.S4.**
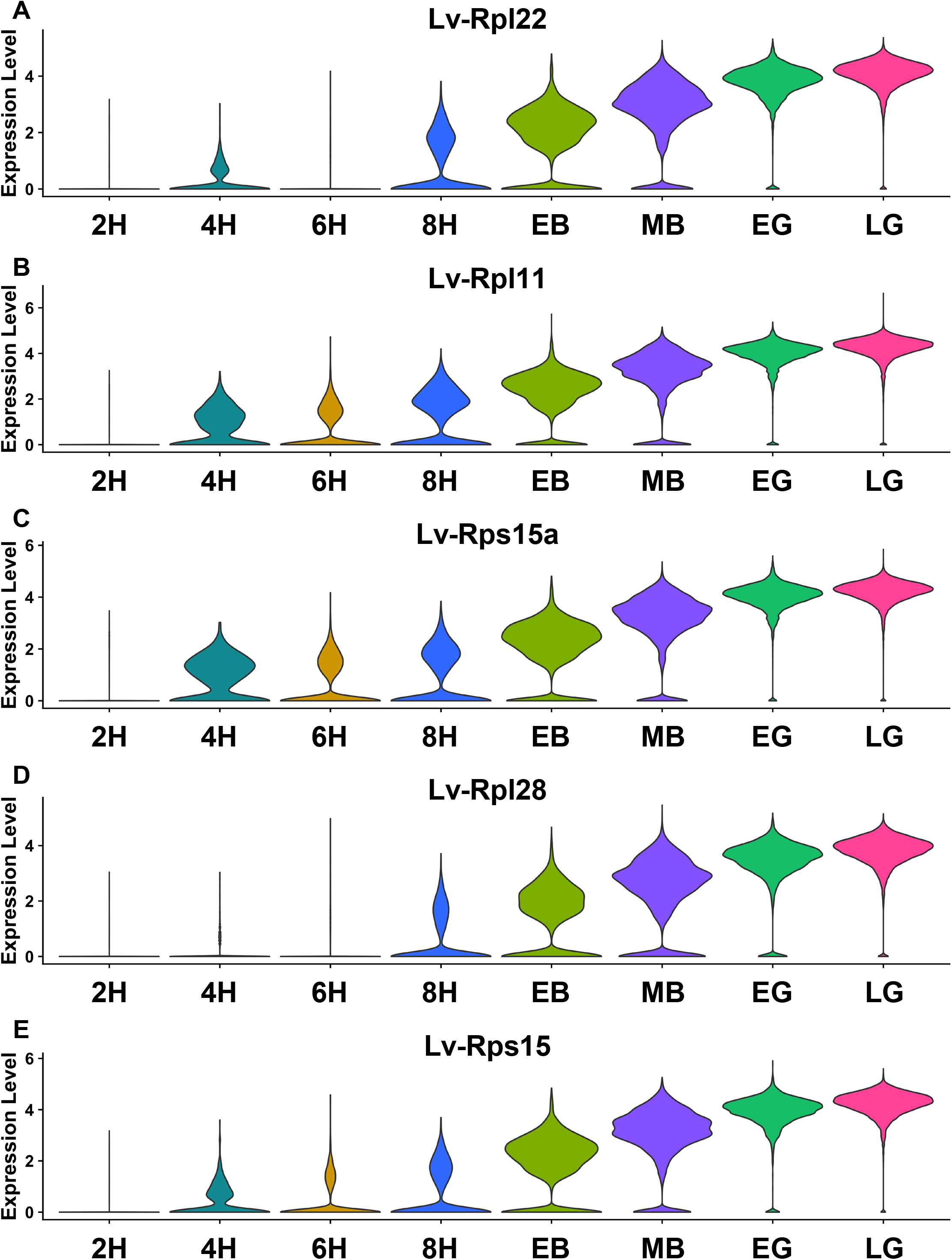
Ribosomal gene expression in the *Lv* dataset. Violin plots showing ribosomal gene expression throughout Lv development

**Fig.S5.**
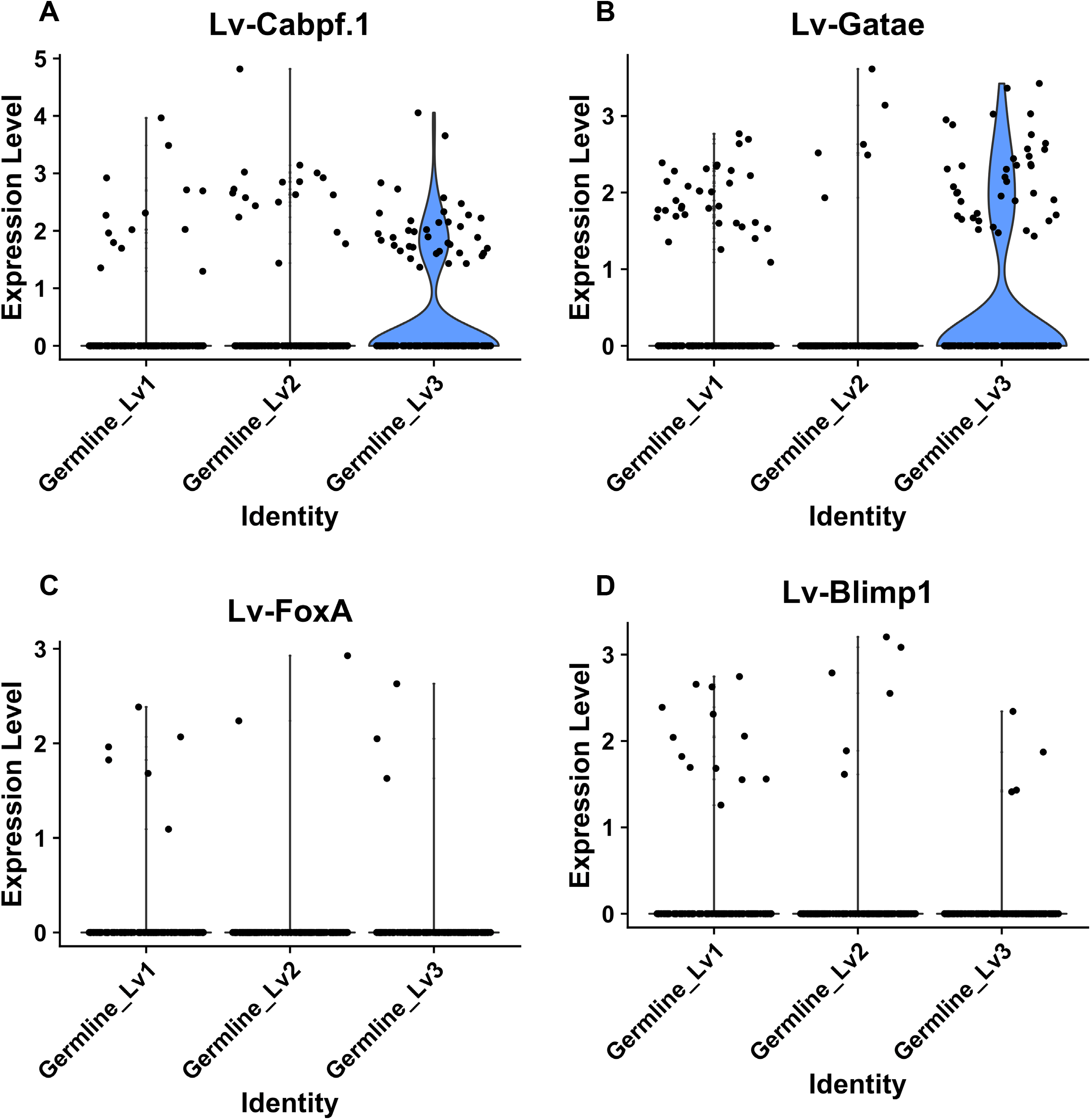
Endodermal gene expression in the germline subpopulations of *Lv* dataset. Violin plots showing endodermal gene expression in Germline_Lv1 (Chatfield *et al*.), Germline_Lv2 (green) and Germline_Lv3 (blue).

**Fig.S6.**
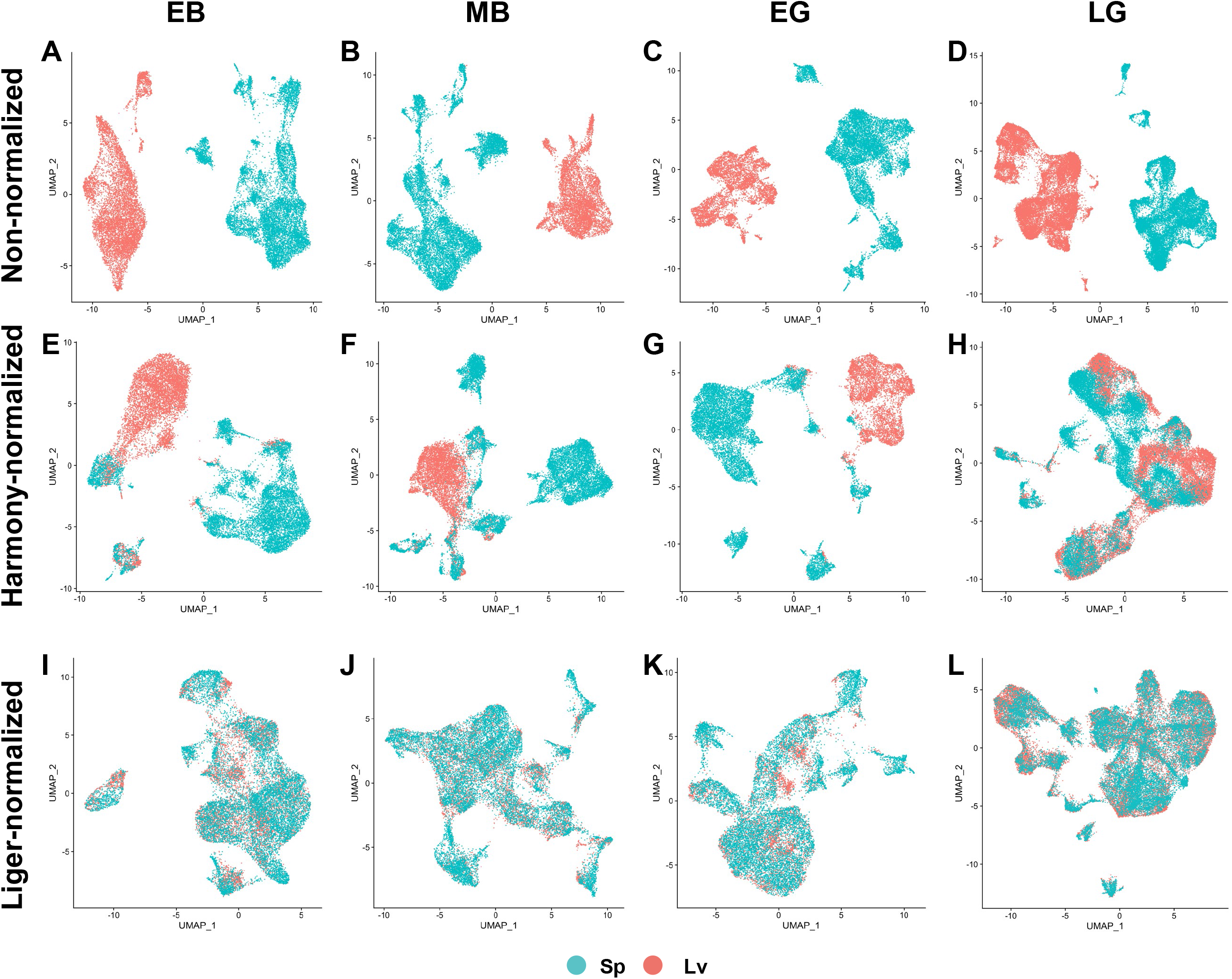
UMAP plot analysis of non-normalized, Harmony-normalized, and Liger-normalized *Lv-Sp* dataset. (A–L) Non-normalized (A–D), Harmony-normalized, and Liger-normalized Lv-Sp datasets are presented. EB (A, E and I), MB (B, F and J), EG (C, G and K) and LG embryo (D, H and L) were integrated individually. Cells are color-coded by the species (*Lv*: red; *Sp*: blue)

**Fig.S7.**
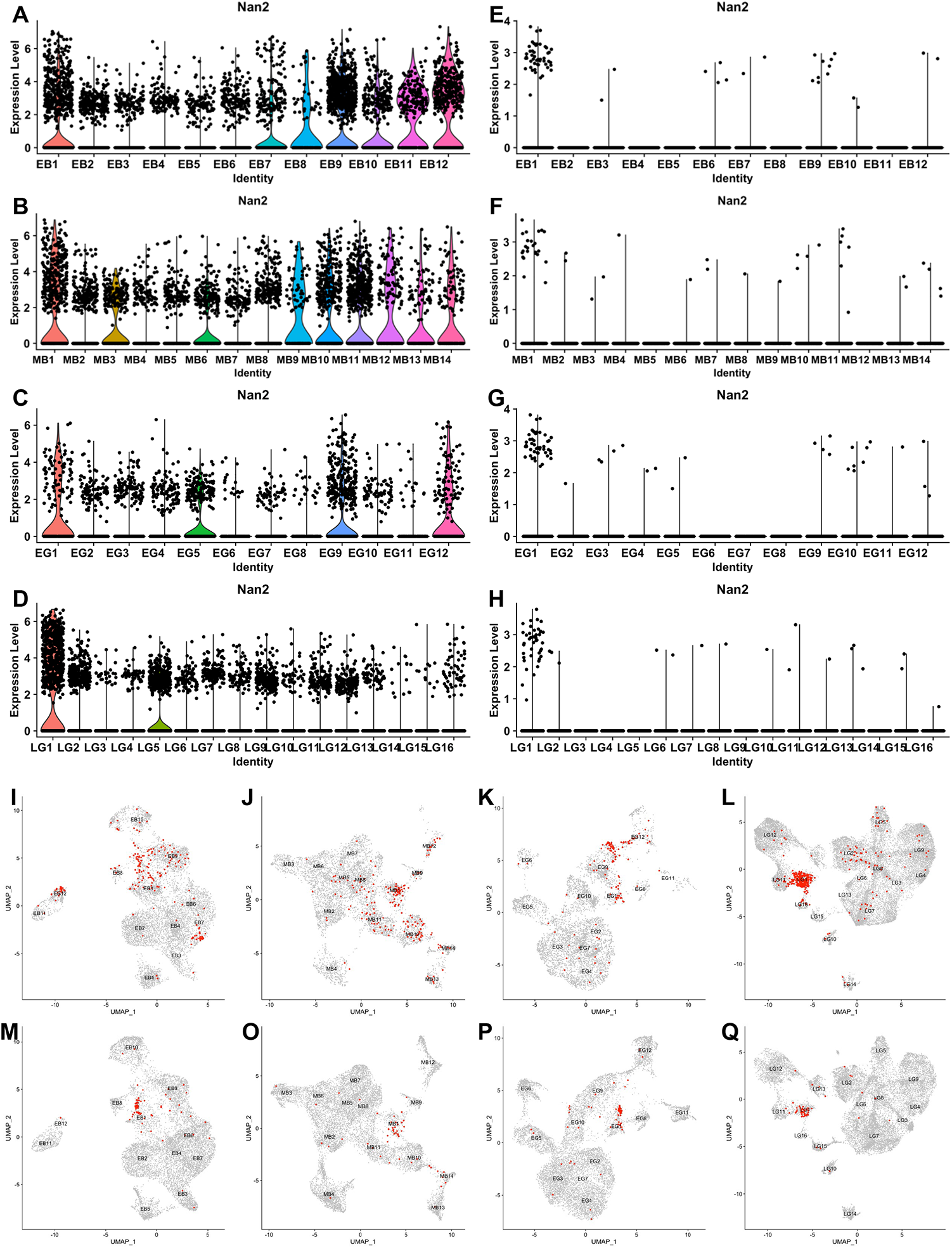
Nanos2 expression and distribution of Nanos2-positive cells in *Lv-Sp* datasets. **(A–H)** Violin plots showing Nanos2 expression in *Lv* (A–D) and *Sp* embryos (E– H). **(I–Q)** Nanos2-positive cells in *Lv* (I–L) and *Sp* embryos (M–Q) are presented as red dots in UMAP plots.

**Fig. S8.**
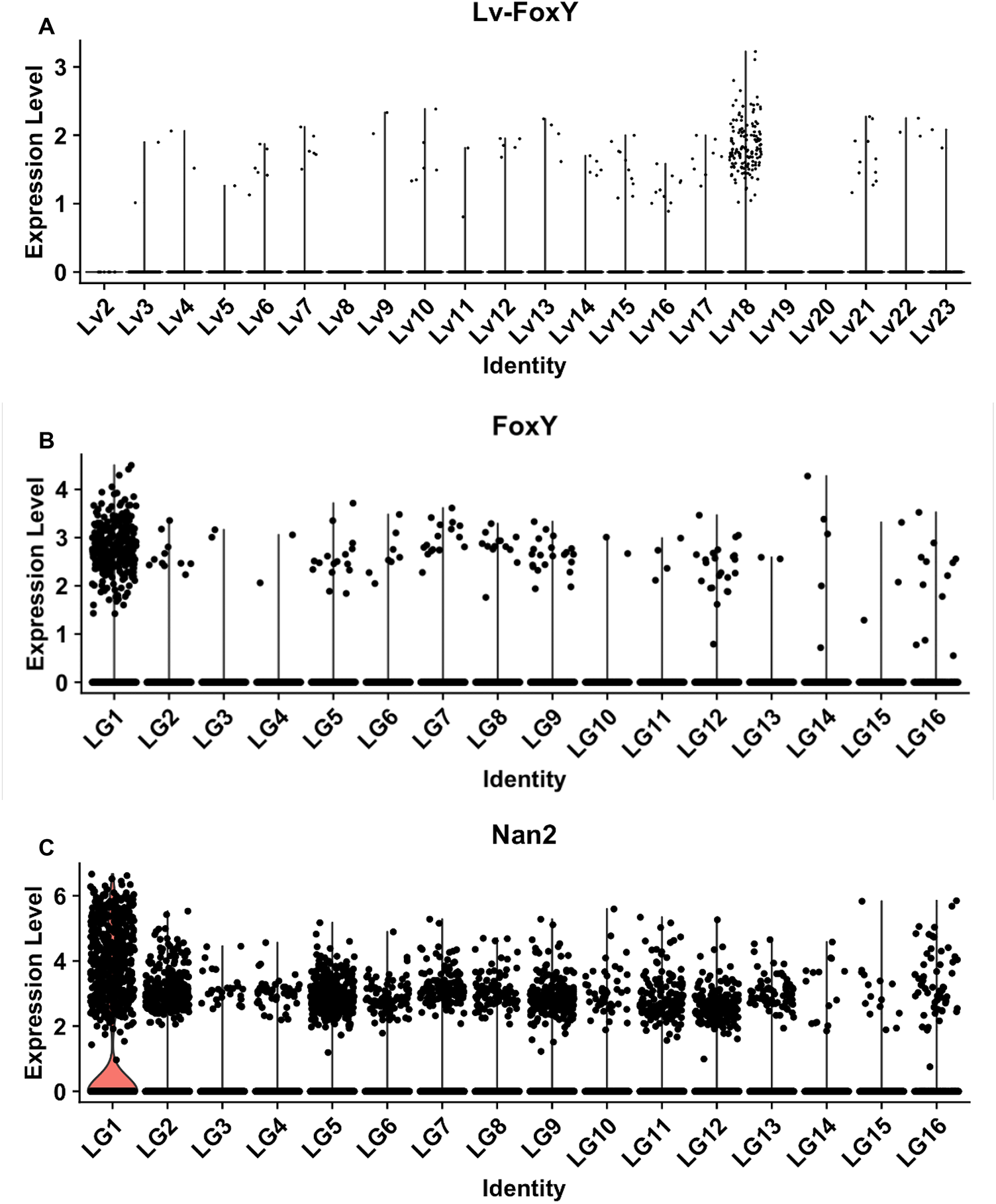
Nanos2 and FoxY expression. (A–C) Violin plots showing *Lv* FoxY expression in the *Lv* dataset (A), FoxY expression in the LG embryos of the Lv-Sp dataset (B) and Nanos2 expression in the LG embryos of the Lv-Sp dataset (C). *Lv* FoxY was enriched in Lv18 mesodermal cells but not in Lv23 Germline cells. On the other hand, both FoxY and Nanos2 were enriched in the LG1.

**Fig. S9.**
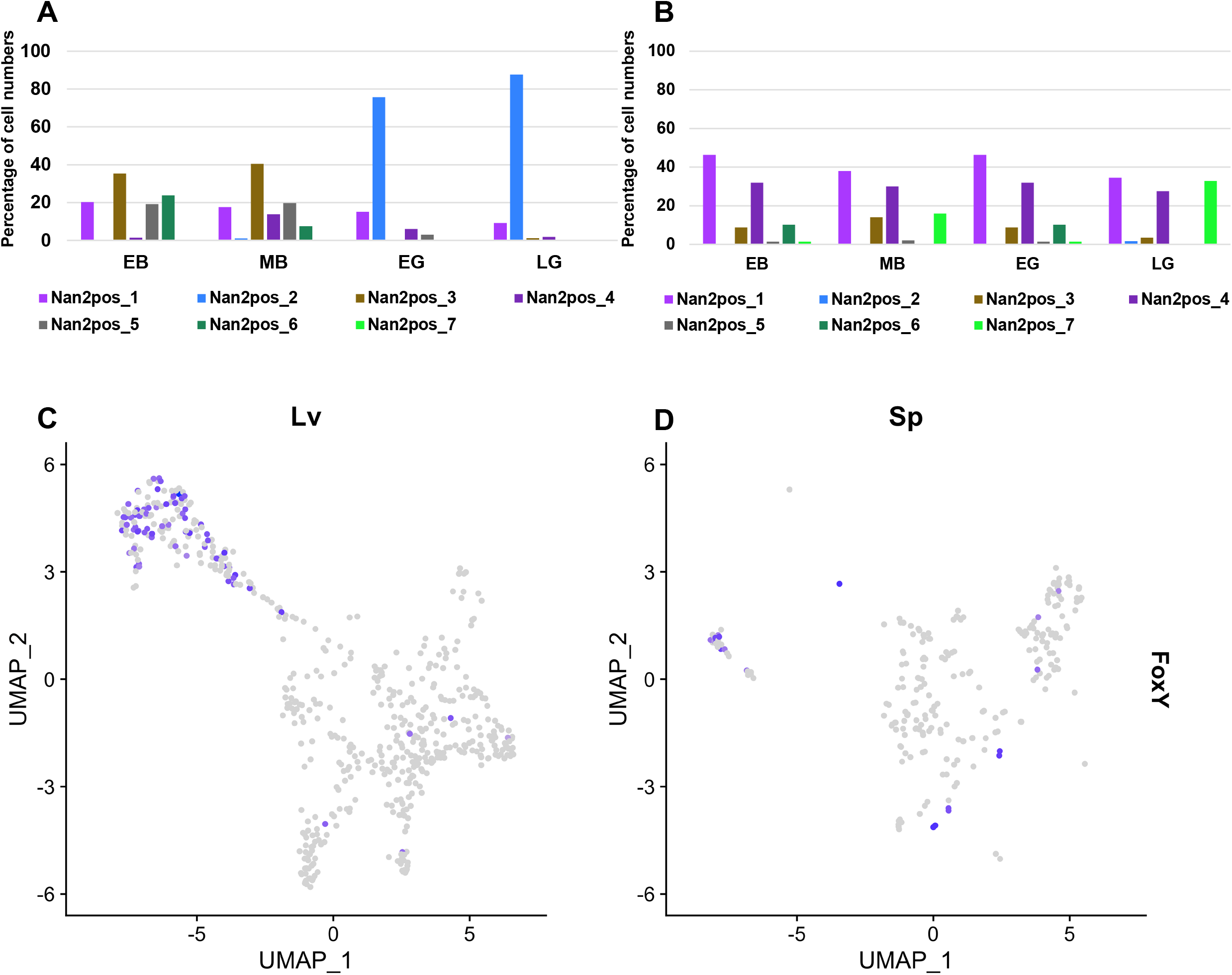
Nanos2 positive subpopulations in the Lv-Sp dataset. The proportion of each Nanos2 positive subpopulation in Lv (A) and *Sp* (B) for each developmental time point. Expression of FoxY in these Nanos2 positive subpopulations in *Lv* (C), and *Sp* (D).

**Fig.S10.**
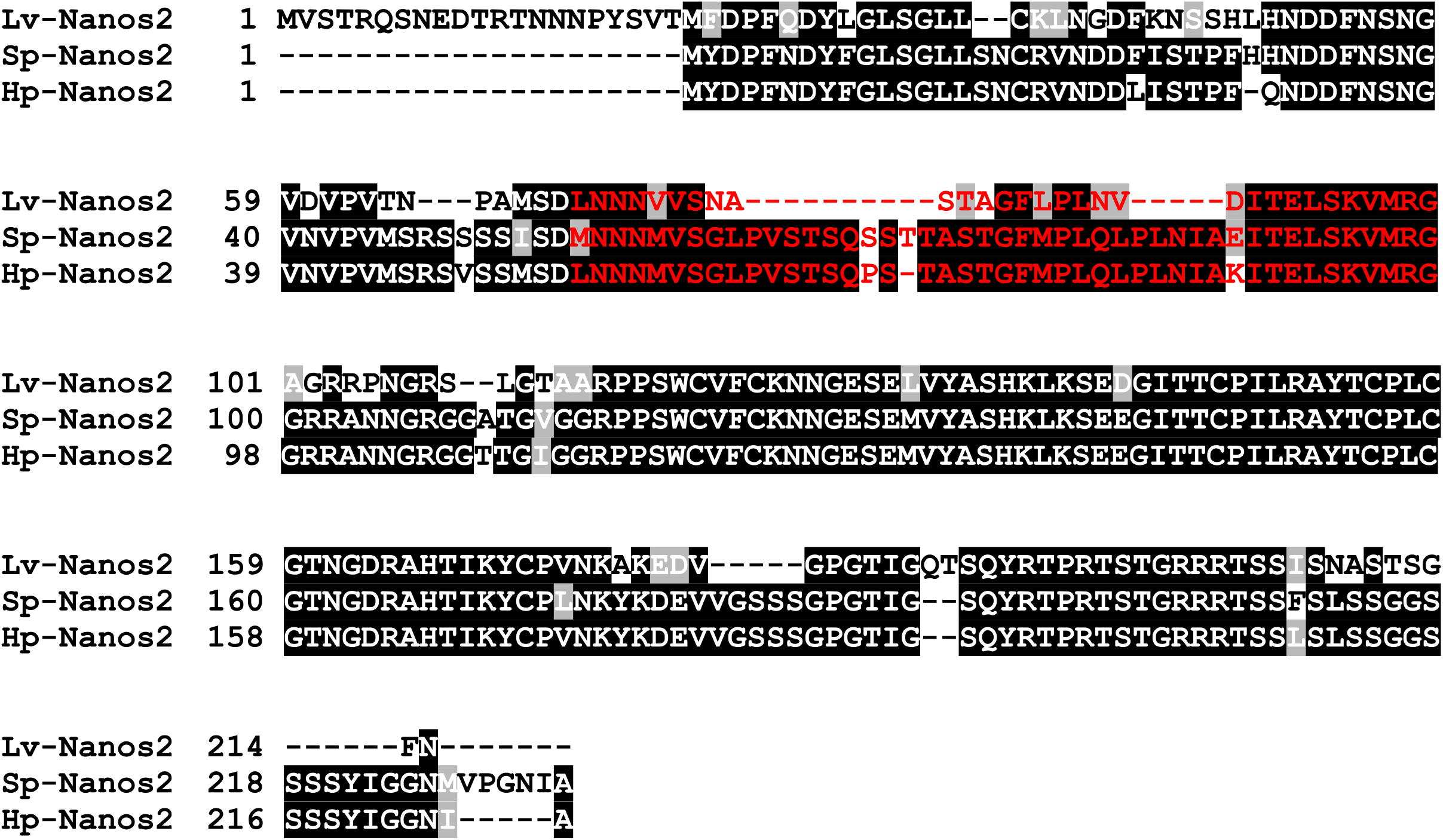
Comparison of Nanos2 protein sequence between species. Nanos2 protein sequences from *Lv* (upper), *Sp* (middle) and *Hp* (lower) are shown. The alignment was generated with T-coffee program and shaded using Boxshade server. Identical and similar amino acid residues between species are shown on black and gray backgrounds, respectively. NPDE sequence is highlighted by red characters.

**Fig.S11.**
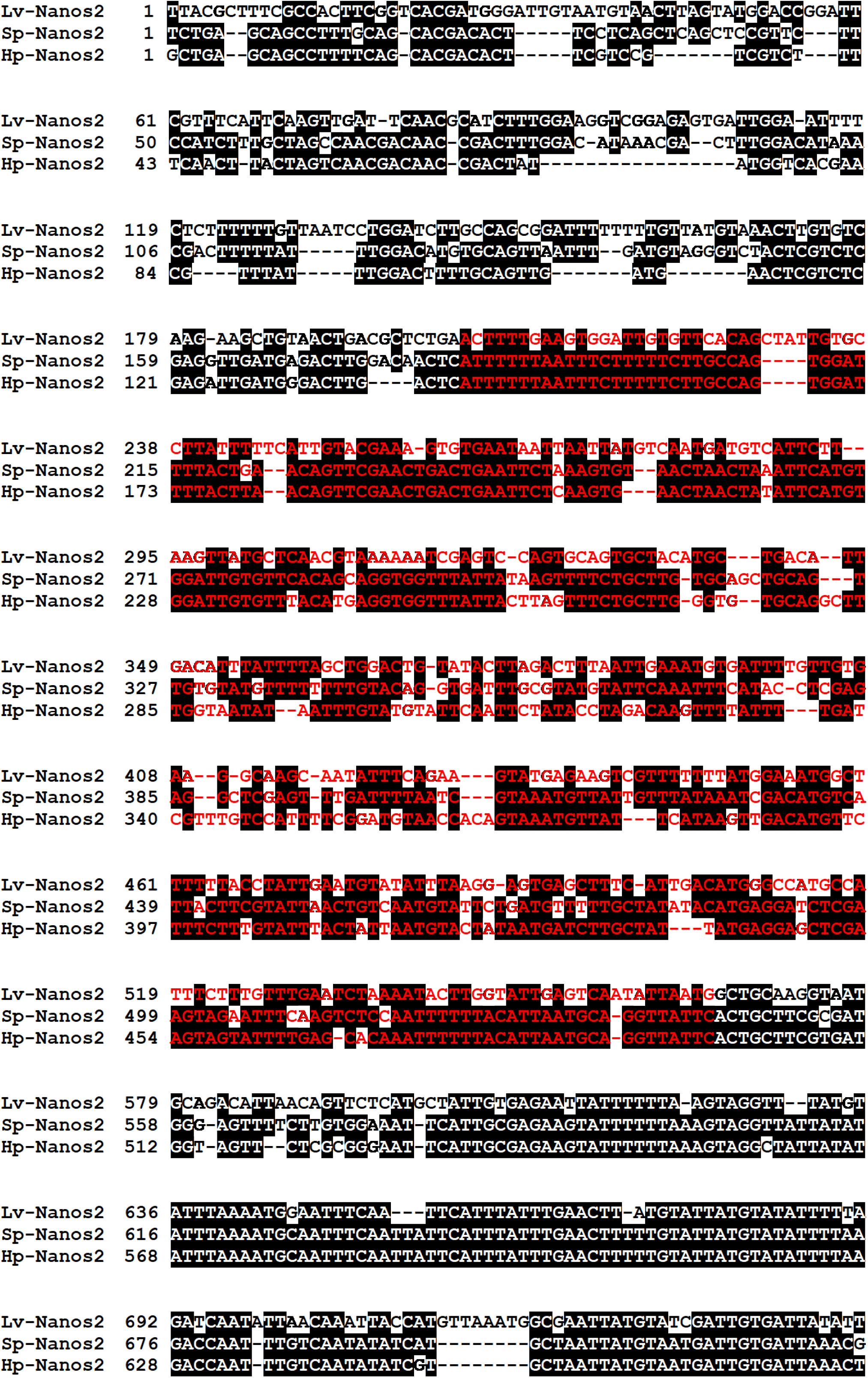
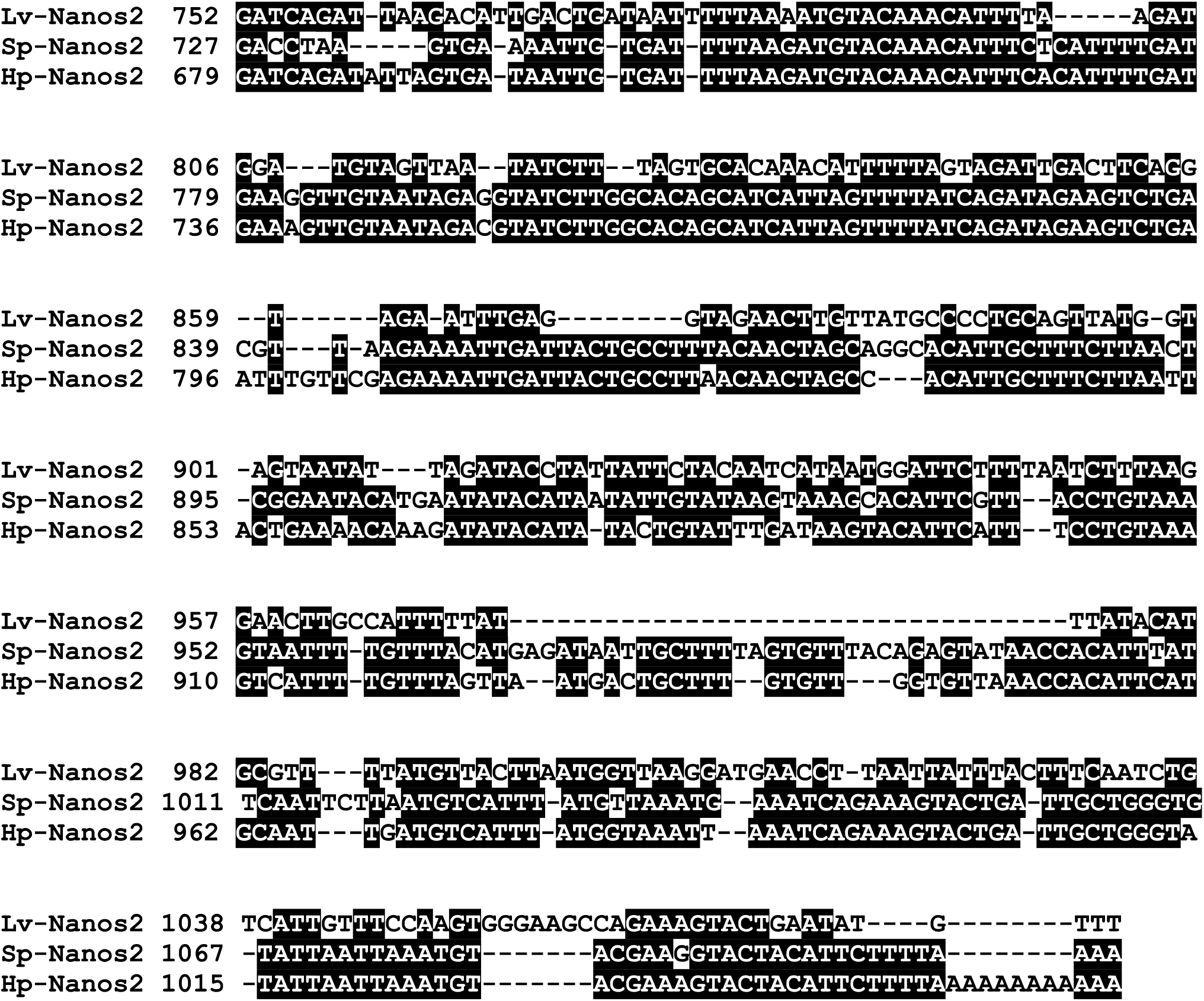
Comparison of Nanos2 3’UTR sequence between species. Nanos2 3’UTR sequences from *Lv* (upper), *Sp* (middle) and *Hp* (lower) are shown. These sequences were compared with T-Coffee with default settings ((Notredame *et al*., 2000); https://tcoffee.crg.eu/apps/tcoffee/index.html) and the result was visualized by BOXSHADE (http://arete.ibb.waw.pl/PL/html/boxshade.html). GNARLE sequence is highlighted by red characters.

**Fig. S12.**
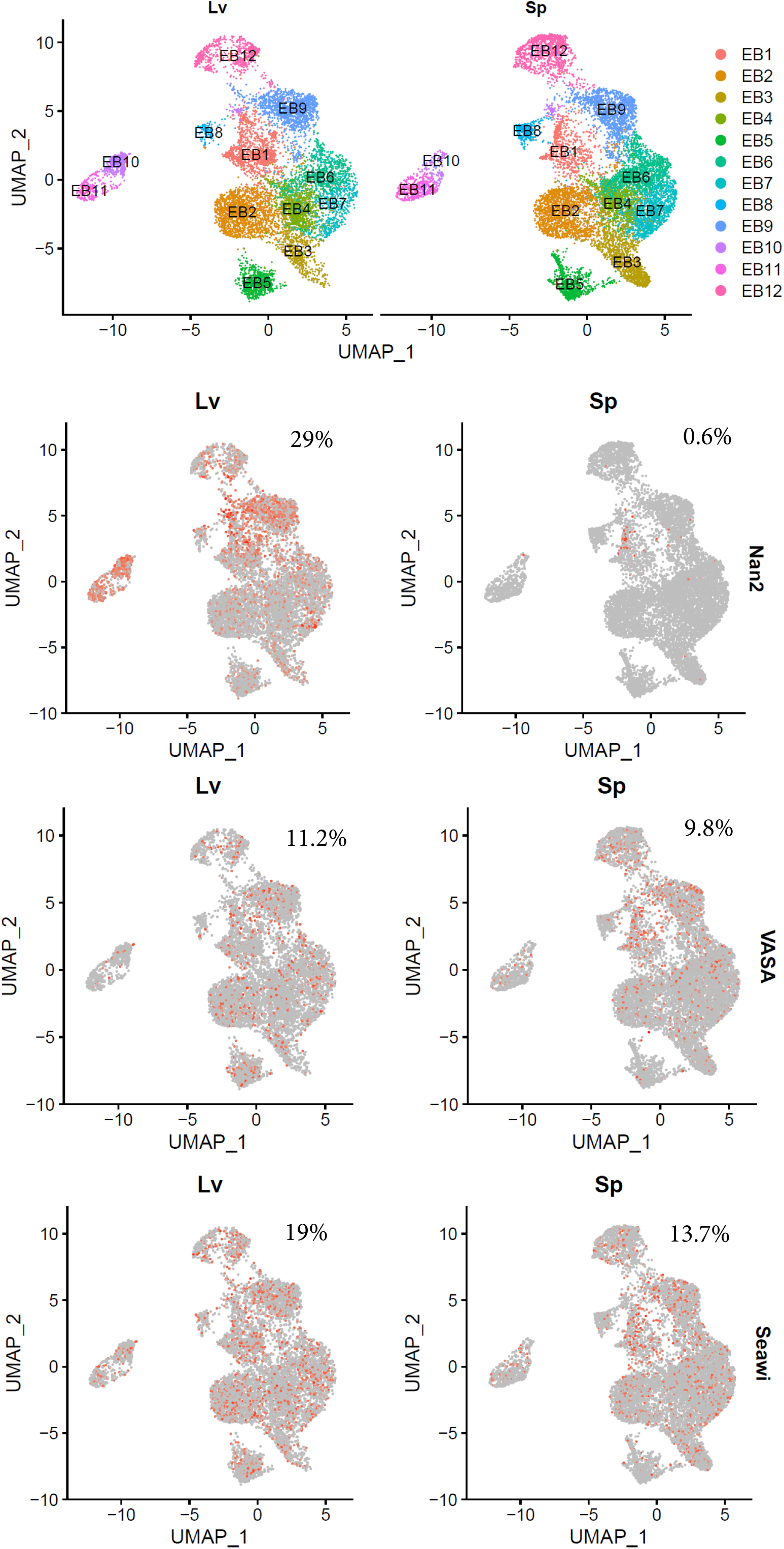
Expression of germline genes in the Lv-Sp dataset at early blastula stage. Umap of the early blastula stage in *Lv* and *Sp* (A), and expression of Nanos2 (B), Vasa (C), and Seawi (D). The number above each plot represents the percentage of cells expressing the corresponding gene.

